# Phosphorus availability as the primary determinant of nutrient limitation in temperate biodiverse herbaceous vegetation

**DOI:** 10.1101/2024.10.18.619065

**Authors:** Kevin Van Sundert, Rudy van Diggelen, Jos D’Haese, Camiel J.S. Aggenbach, Enrico Dammers, Sophia Findeisen, Wiktor Kotowski, Lucasz Kozub, Richard Kranenburg, Arrie van der Bij, Willem-Jan Emsens

## Abstract

Understanding whether nitrogen (N), phosphorus (P) or potassium (K) (co)limit productivity across biodiverse herbaceous habitats is crucial to guide management. Therefore, we investigated for 386 plots representing 13 nutrient-limited habitat types across Europe whether community N:P:K stoichiometry and limitation types differ along wide-ranging gradients in soil development, moisture and pH. Results indicate P/P+N as frequent as N limitation. K/K+N limitation occurs not where K availability is minimal, but in species-impoverished habitats with excess N and P. Overall, P emerges as primary driver of stoichiometry, strongly driven by the environment: at optimal pH of 6, N:P and P/P+N limitation are minimal and N limitation maximal, despite also good conditions for N availability. At pH<5 and >7, N:P is high and P/P+N limitation common. Our findings emphasize soil pH control on nutrient limitation through influence on P. Studies reporting widespread K (co)limitation in temperate herbaceous vegetation likely sampled anthropogenically P/N-enriched communities.

---

Plant species evolved adaptations to survive, grow and reproduce within environmental conditions that they can physiologically tolerate. At the extreme end of such environmental conditions (e.g. waterlogging^1^, severe water shortage, extreme acidity or alkalinity), the environment becomes a strong abiotic filter that directly constrains the plant community assembly. In less extreme environments without frequent disturbance, community assembly is determined by competition among plants for resources. Where productivity is high and nutrients are ample, competition is strongest for light. Where productivity and nutrient availability are lower, plants compete for these nutrients. It is in such nutrient-limited but non-extreme environments that highly diverse plant communities with rare species are found^2^. Understanding the nature of limitations and competition for nutrients therefore contributes to the conservation and restoration of biodiversity.

From an ecosystem ecological perspective, ‘limitation’ of a process (e.g. productivity) by a resource such as nutrients is defined as ‘demonstrable when a substantial addition of a particular element increases the rate or endpoint of that process’^3–5^. Across terrestrial natural and semi-natural ecosystems globally, limitation by nutrients such as nitrogen (N), phosphorus (P) and/or potassium (K) is the most widespread type of limitation^6,7^. Biomass production of nutrient-limited vegetation is not determined by the total available amount of all these soil nutrients^8^. Instead, some nutrients can be present in an oversupply, while at the same time another nutrient or set of nutrients has an availability at such level that it limits plant growth. The productivity of vegetation only increases substantially when such limiting nutrient(s) are added^9^. Accordingly, the traditional approach to determine the limiting nutrient(s) are fertilization experiments^10,11^. In such experiments, single or combined nutrients are added, after which an assessment is made of in which cases an increase in productivity or in other ecosystem processes occurs. Much research has been performed in this way, but this is an invasive and labor intensive approach.

An alternative method to determine nutrient limitation is based on the ratios of the mass-based abundance of N, P and K in vascular plant tissue. Better than absolute nutrient concentrations, ratios express the relative availabilities of these nutrients^5^. Calibrated on fertilization experiments in wetlands, Koerselman and Meuleman (1996)^12^ and Olde Venterink et al. (2003)^13^ defined critical N:P, N:K and K:P ratios that discern N limited versus P/P+N colimited versus K/K+N colimited growth. Further research showed that N:P:K ratios effectively varied more strongly among sites than among species, and that these ratios correlated well with effects of nutrient fertilization^14,15^. Several researchers have applied the use of these critical ratios in studies on herbaceous vegetation such as in wetlands^16–18^, grasslands^18–20^ and forest understories^15^. N:P:K ratios appeared to be good indicators of the type of limitation, provided that nutrients are effectively limiting factors.

Nitrogen is the most commonly limiting nutrient in vegetation globally, and particularly in temperate and cooler climate zones^3,21^. The availability of N is strongly controlled by microbial processes, such as biological dinitrogen fixation that provides new input of N into the ecosystem^22^, mineralization that releases plant-available N from unavailable N in organic matter, nitrification that converts ammonium to nitrate, and denitrification that removes N from the system in gaseous form^23^. As a result, N availability typically increases as time has passed for N to accumulate in ecosystems through fixation^24^. N availability generally decreases with increasing soil moisture, as particularly wet and anaerobic conditions preclude high microbial activity^25,26^. Additional to that, microbial activity is strongly affected by soil pH, with an optimum between 5.5-6 and 7^27^. Finally, N availability has strongly been influenced by anthropogenic activities. In particular N deposition has had profound effects on the plant-availability of N relative to that of other elements such as P and K, to such extent that naturally N limited vegetation can shift to (co)limitation by P^28^ or other macro- and micronutrients.

After N, P is the second most commonly limiting nutrient^29^. Compared to N, P availability is less affected by microbial processes but more by physicochemical processes^30^. P has no gaseous form which can be drawn upon from the atmosphere (although some P deposition does occur^31^) and P-compounds are poorly soluble. P is therefore much less mobile than N and P-availability declines with time, when the topsoil gets increasingly depleted of P^32^. Soil P content can increase in sites where influx of groundwater occurs^33^, but the effect on P-availability depends on pH. Indeed, P availability is strongly determined by soil acidity, with optimal conditions occurring at around pH = 6-7.5: at lower pH, P adsorbs to aluminum and iron oxides^30,34^, whereas at higher pH calcium binds P and stimulates its adsorption to soil minerals, making it less accessible for plant uptake^35–37^. Low pH further correlates with low P for rainwater-fed and thus acidic peatlands (i.e. bogs^38^), which exhibit near-zero P input.

Besides N and P, K is a third macronutrient that can limit vegetation processes^39^. The in situ source of available K to plants in terrestrial ecosystems is the primary weathering of silicates^40^. Apart from atmospheric deposition^31^ and surface water influence, important pathways for K to leave or enter ecosystems pass through soil and groundwater, since K-salts are easily water-soluble - more so than P containing compounds^41^. K is easily leached out (its retention depending on soil properties such as cation exchange capacity^40^), and also K influx in base rich groundwater is an important mechanism influencing local soil concentrations of available K^26^. Relative to plant demand, K supply is generally more abundant (and less often limiting) than that of N and P^29^. K availability can be low, however, in soils with particularly low cation exchange capacity (CEC)^40^, low soil moisture inhibiting its diffusion^42^, or under acidic circumstances with desorption of K from exchange sites and subsequent leaching^43^.

While these overall described roles of soil age and environmental gradients such as soil moisture and pH in influencing N, P and K availabilities have been well established, considered over a broad range of habitats it remains unclear which nutrients are most limiting under what environmental conditions. We therefore tested hypotheses on relative availabilities and limitations of N, P and K against vegetation stoichiometric and environmental data from 386 herbaceous biodiverse plant communities across northwest and central Europe. Community composition in these habitats was *a priori* expected to be driven by competition for soil resources rather than light given the relatively low productivity (e.g. standing biomass^44^ mostly < 6000 kg ha^−1^, Table S1). The variety of 13 studied habitat types, with uniquely to our study, particularly long gradients in stages of soil development, soil moisture and pH allowed us to assess the general applicability of theories on nutrient limitations and relative availabilities across temperate and herbaceous nutrient-limited vegetation. Specifically, we expect:

the availability of N to increase, and that of P to decrease as soil development progresses. Consequently, N:P and N:K ratios, as well as P/P+N and K/K+N limitation increase and N limitation decreases with succession-related soil N buildup.

the availability of N to decrease with increasing soil moisture (following slower decomposition), and that of P and K to increase with increasing soil moisture (because of higher diffusion and desorption rates). Consequently, N:P and N:K ratios, as well as P/P+N and K/K+N limitation decrease and N limitation increases under elevated soil moisture.

a slightly acidic to circumneutral pH to be optimal for the availability of both N and P, and K availability to increase particularly strongly with pH. Consequently, we expect N:K and the occurrence of K/K+N limitation to decrease and K:P to increase with pH.

plant communities on locations with higher N deposition to have elevated N:P and N:K ratios, resulting in shifts from N limitation to P/P+N and K/K+N limitation.

finally, anticipating N limitation to be the ‘default’ limitation type in the temperate region^45^, but also high spatial variation in available N through the gradients mentioned above as well as anthropogenic N deposition^46^, we hypothesize the availability of N to be the strongest driver of stoichiometric ratios and limitation types. After N, we expect P availability to be a stronger driver than K, as we foresee only rare occurrence of K (co)limitation *a priori*^29^.

## Results

### Nutrient limitation among habitat types

Following within- and among-habitat variation in N:P:K ratios and deviations from critical ratios (Figs. 1a and 2a, Text S1), also nutrient limitation types varied within and among habitat types (Figs. 1a and 2b). Overall, 178 sites were N limited, 165 P/P+N limited, 28 were K/K+N limited and 15 did not fit into any of the categories. K (co)limitation appeared particularly in moist grasslands (*n* = 18 out of 65 sites) and dry grasslands (4 out of 21 sites), and in few fens (*n* = 2), fen meadows (*n* = 1), *Nardus* grasslands (*n* = 2) and dry heathlands (*n* = 1). Focusing on the more common N and P (co)limitations, proportions of N-versus P/P+N limitation in vascular plants differed significantly among habitat types (χ²_12_ = 190.33, *P* < 0.001; Fig. 2b). Four groups could be discerned amongst the habitat types: fens, dry dunes and wet eutrophic marshes, which are mostly N limited; wet dune slacks, dry calcareous grasslands and *Nardus* grasslands, with approximately equal occurrence of both types of limitation; calcareous springs, bogs, dry and wet heathlands that are predominantly P/P+N limited; and the more species-impoverished moist and dry grasslands, which are predominantly N limited but where K/K+N limitation also commonly occurs.

**Fig. 1.**
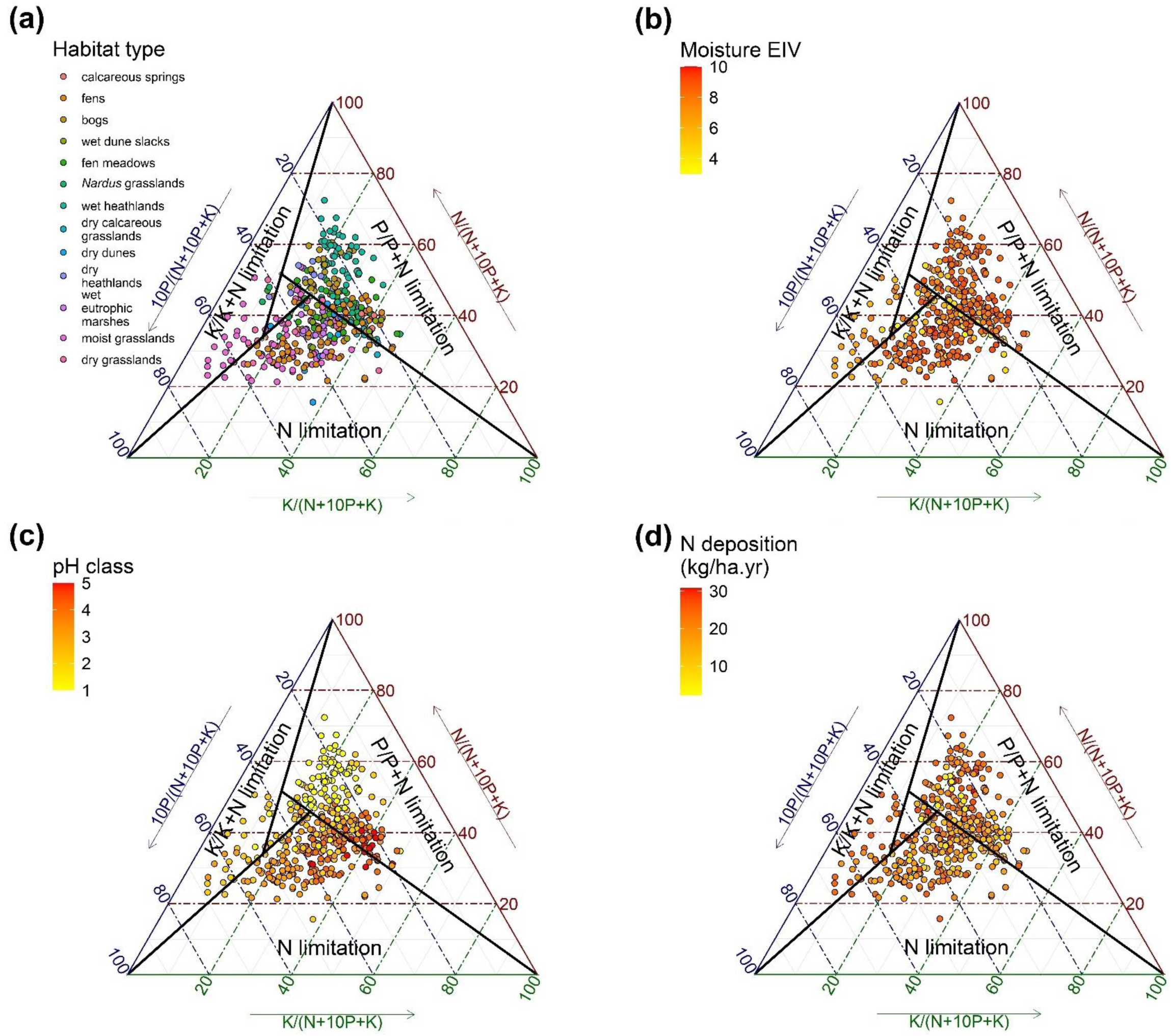
Distribution of sampling points by relative vegetation N, P and K concentrations and limitation types. Point colors indicate stratification by **a**, habitat type, **b**, Ellenberg Indicator Value (EIV) for soil moisture, **c**, pH class and **d**, N deposition. For sites in the middle area of the ternary diagrams, the type of nutrient limitation could not be assigned based on N:P:K ratios. Phosphorus concentrations were multiplied by 10 to improve the spread of data points in the visualization. pH classes are: 1 = acidic, 2 = moderately acidic, 3 = weakly acidic, 4 = circumneutral, 5 = alkaline.

**Fig. 2.**
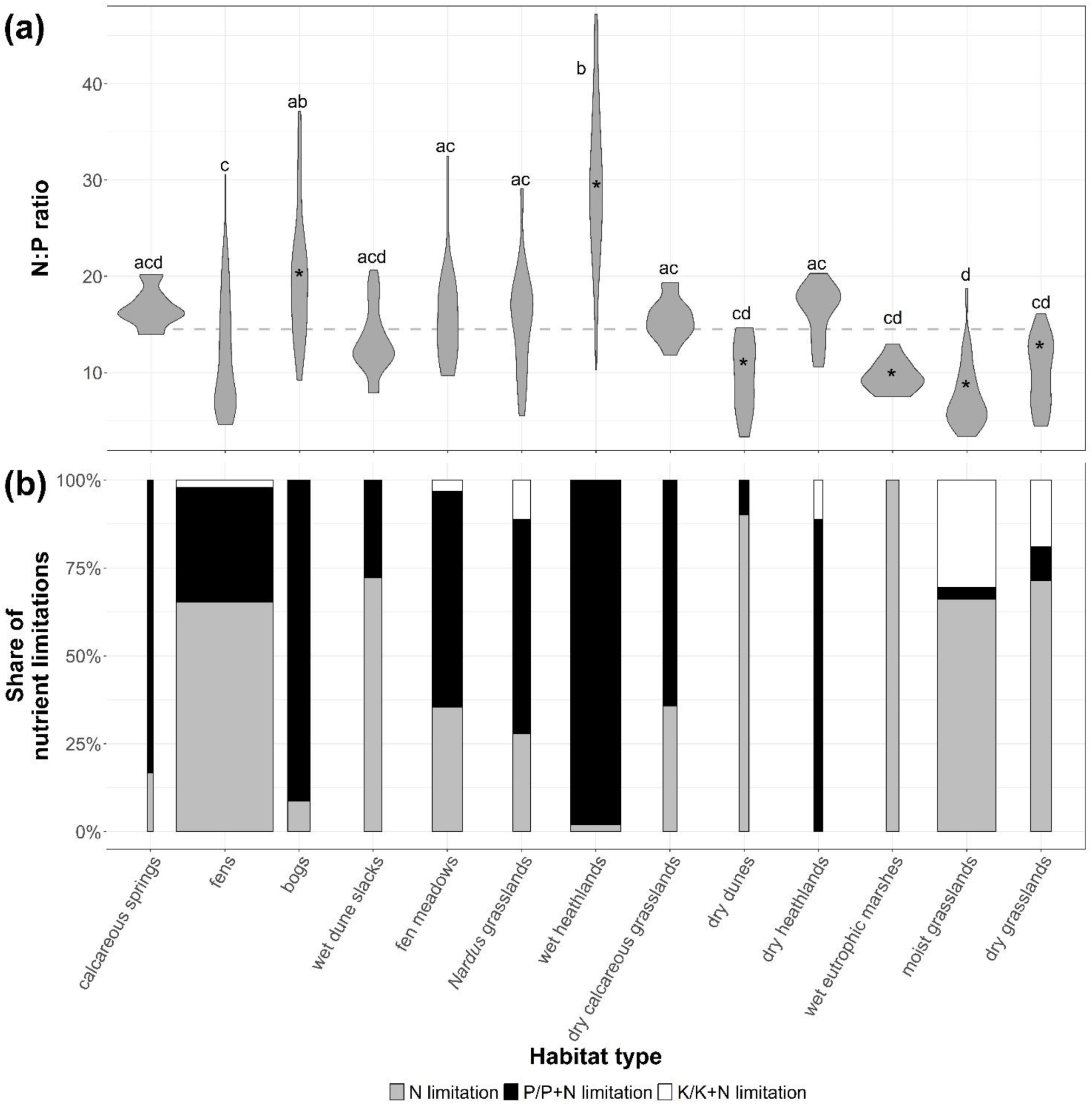
Patterns of vegetation N:P stoichiometry (g g^−1^) and derived nutrient limitation types across habitat types. **a**, the N:P ratio, with critical ratio of 14.5^13^ indicated through a dashed line. **b**, nutrient (co)limitation types. In **(a)**, values significantly differing from the critical N:P ratio were marked with ‘*’, values significantly different among habitat types were indicated trough different letters. Widths of bars in **(b)** were scaled by the number of sites (calcareous springs: *n* = 6; fens: *n* = 99; bogs: *n* = 26; wet dune slacks: *n* = 19; fen meadows: *n* = 32; *Nardus* grasslands: *n* = 18; wet heathlands: *n* = 51; dry calcareous grasslands: *n* = 14; dry dunes: *n* = 11; dry heathlands: *n* = 11; wet eutrophic marshes: *n* = 13; moist grasslands: *n* = 65; dry grasslands: *n* = 21).

### Nutrient limitation and environmental gradients

Nutrient stoichiometry and limitation types clearly related to soil pH, but relationships with soil moisture and atmospheric N deposition seemed less apparent from the visual spread of nutrient limitations along ecological gradients (Fig. 1b-d). For soil moisture, formal analysis revealed that over the entire dataset, N:P and therewith P/P+N limitation did significantly increase with an increasing moisture EIV, but with much variation remaining unexplained (*R²*_m_ ≤ 0.03). Also vegetation K increased, and N:K as well as K/K+N limitation accordingly decreased with increasing moisture, again with low *R²*_m_ ≤ 0.02. N limitation was unrelated to soil moisture (Table 1).

**Table 1.**
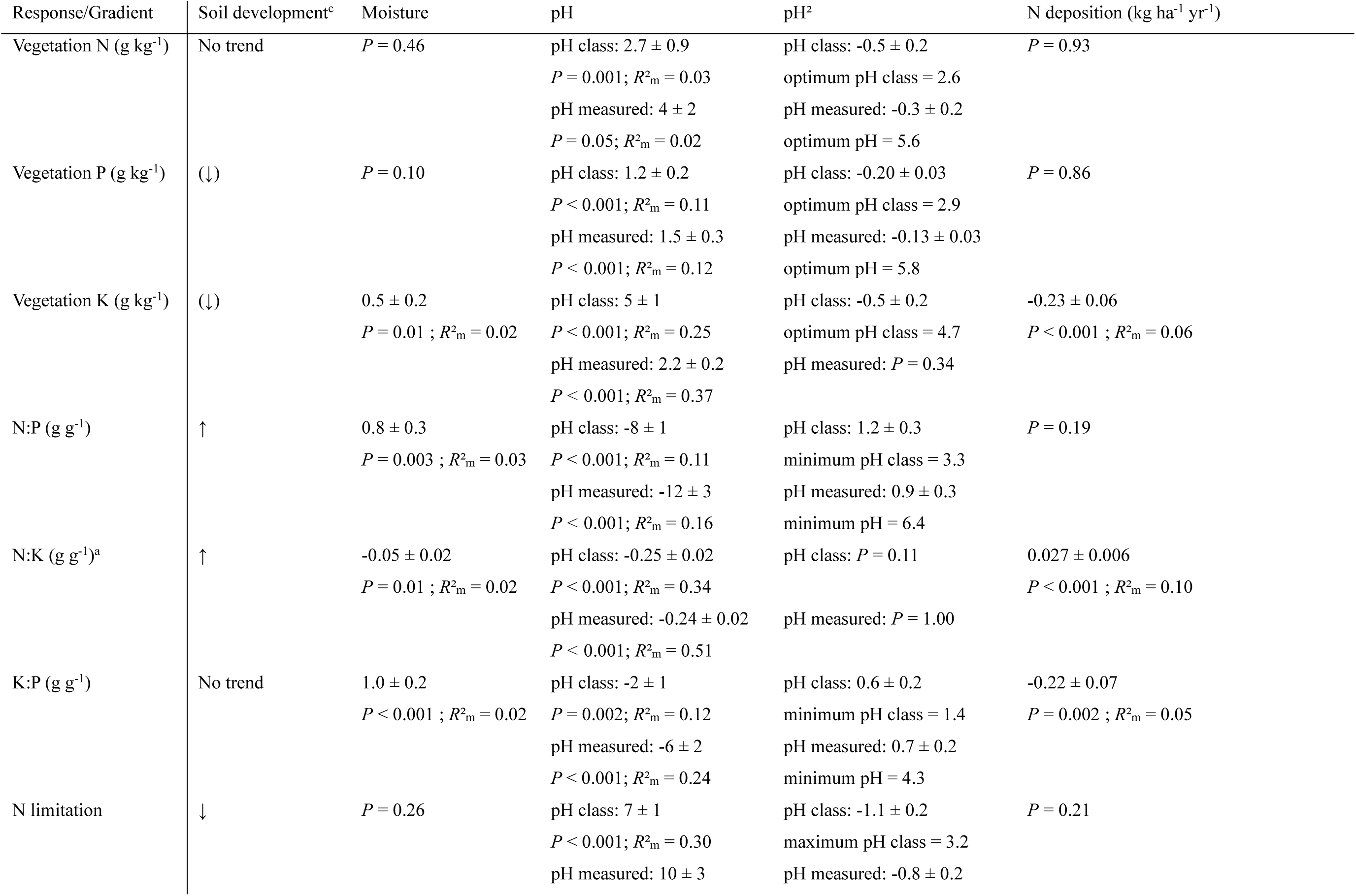
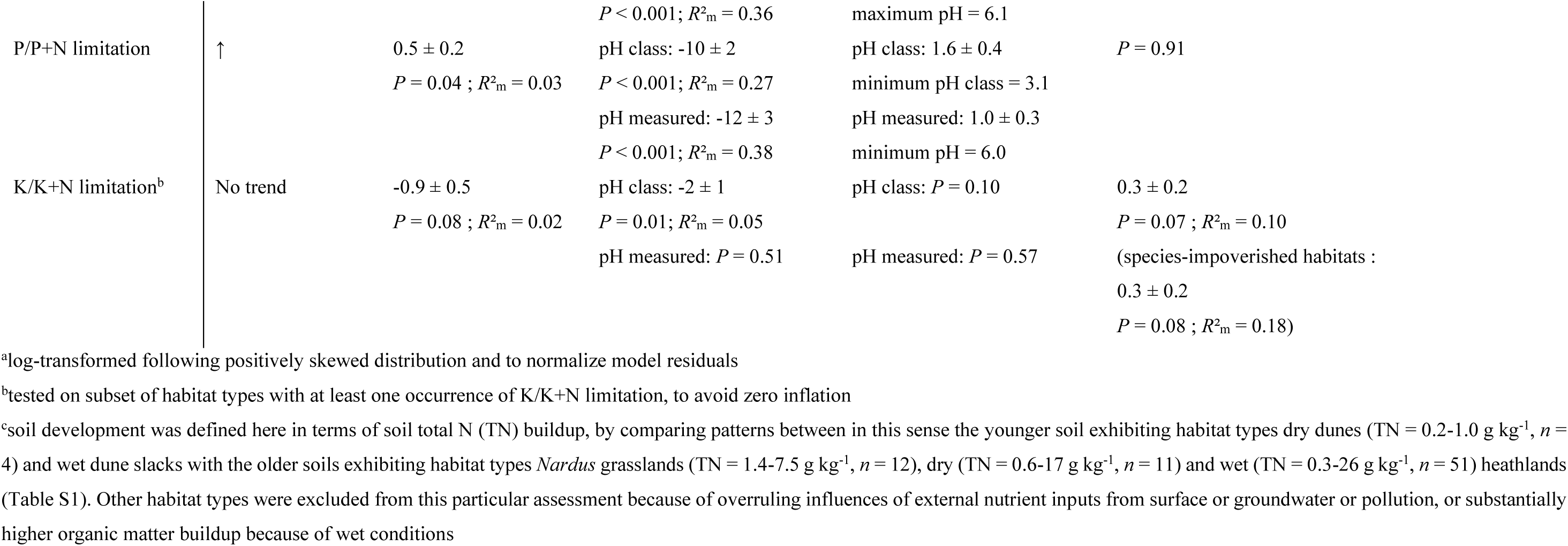
Associations of vegetation N, P and K concentrations, stoichiometry and limitation types with soil development and environmental gradients.

Visual presentation suggests soil pH as a key driver of vegetation stoichiometries and limitation types (Fig. 1c). P concentrations relative to N appear to increase with pH up to the circumneutral pH class 4, above which relative P concentrations drop again. Consequently, P (co)limitation occurs where soils are either acidic, or strongly alkaline, whereas N limitation occurs on soils with intermediate pH. Formal analysis here confirms the major impact of soil pH on vegetation nutrient concentrations, stoichiometry and limitation (Table 1). N and P concentrations exhibited optima at around pH = 5.6 (or pH class = 2.6) and pH = 5.8 (or pH class = 2.9), respectively, with P more strongly pH-affected than N (e.g. *R²*_m_ ≥ 0.11 for P and *R²*_m_ ≤ 0.03 for N). This resulted in a minimal N:P at pH = 6.4 (or pH class = 3.3; Fig. 3a), minimal P/P+N limitation at pH = 6.0 (or pH class = 3.1), and maximal N limitation at pH = 6.1 (or pH class = 3.1; Fig. 3b). Vegetation K was strongly and positively related to soil pH (*R²*_m_ = 0.25–0.37), resulting in decreasing N:K (*R²*_m_ = 0.34–0.51; Extended Data Fig. 3c-d), and (above a minimum at low pH) increasing K:P ratios with increasing pH (*R²*_m_ = 0.12–0.24; Extended Data Fig. 3e-f). However, a consequently expected negative relationship between pH and K/K+N limitation was comparatively weak when using classified pH (*R²*_m_ = 0.05) and absent when using measured pH (Table 1).

**Fig. 3.**
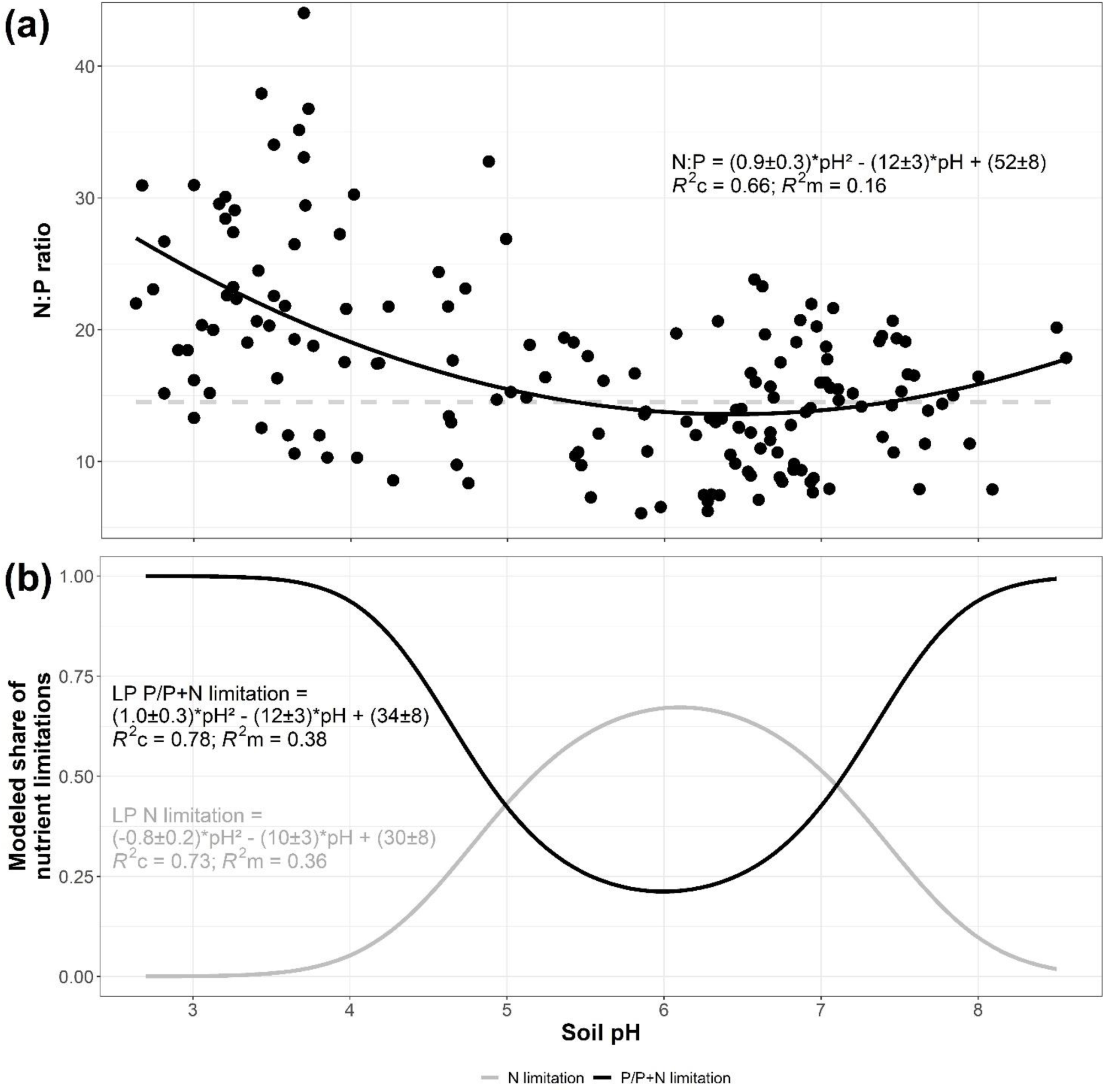
Associations of vegetation N:P stoichiometry (g g^−1^) and probabilities of N versus P (co)limitation with soil pH, measured *in situ* or in water. **a,** the N:P ratio with a minimum around pH = 6.4, and the critical value of 14.5^13^ indicated through a dashed line. **b**, share of P/P+N limitation with a minimum around pH = 6.0, and share of N limitation with a maximum around pH = 6.1. LP in the equations refers to linear predictors. The probability of a nutrient limitation type = exp(LP)/(1+exp(LP)).

Across our dataset, atmospheric N deposition in isolation did not occur as a primary driver of the vegetation’s elemental concentrations, stoichiometry and limitation type (Fig. 1), although in more eutrophic and species-poorer habitat types (wet eutrophic marshes, moist and dry grasslands), it did contribute to overcoming N limitation, inducing K/K+N limitation instead (Table 1). When accounting for N deposition in multiple regression models incorporating the influence of both soil moisture and pH, N deposition did neither modify the (remaining – Extended Data Table 2) dominant role of pH, but it did add explanatory power (*R²*_m_ without N deposition = 0.22; *R²*_m_ with N deposition = 0.32) in particular in describing N:P when measured pH was used (Extended Data Table 3). When testing for the influence of soil moisture, pH and N deposition concurrently, N deposition thus contributes to elevated N:P, partly dependent on pH.

### Which nutrient drives the limitation?

P/P+N limitation occurred where vegetation P concentrations were minimal, as opposed to N limitation which often occurred where vegetation N concentrations were not the lowest (Fig. 4d). For K/K+N limitation, an intermediate situation appeared where K concentrations were on average significantly lower where this type of limitation occurred, but the habitat types with the lowest K concentrations in particular (bogs: 5 ± 1 g kg^−1^, wet heathlands: 5.5 ± 0.9 g kg^−1^, dry heathlands: 5 ± 1 g kg^−1^), still rarely exhibited the K/K+N limitation; instead, this type of (co)limitation was more common in dry and particularly moist grasslands (Fig. 2b) with intermediate K concentrations (dry grasslands: 9 ± 1 g kg^−1^, moist grasslands: 10.4 ± 0.7 g kg^−1^) combined with high N (dry grasslands: 13 ± 1 g kg^−1^, moist grasslands: 16.1 ± 0.6 g kg^−1^) and especially P (dry grasslands: 1.4 ± 0.2 g kg^−1^, moist grasslands: 2.3 ± 0.1 g kg^−1^). Combining boxplots of vegetation N, P and K concentrations also clearly indicates that K/K+N limitation occurred where K was relatively low while at the same time, N and P were significantly higher than for both N and P/P+N limited vegetation (Fig. 4d).

**Fig. 4.**
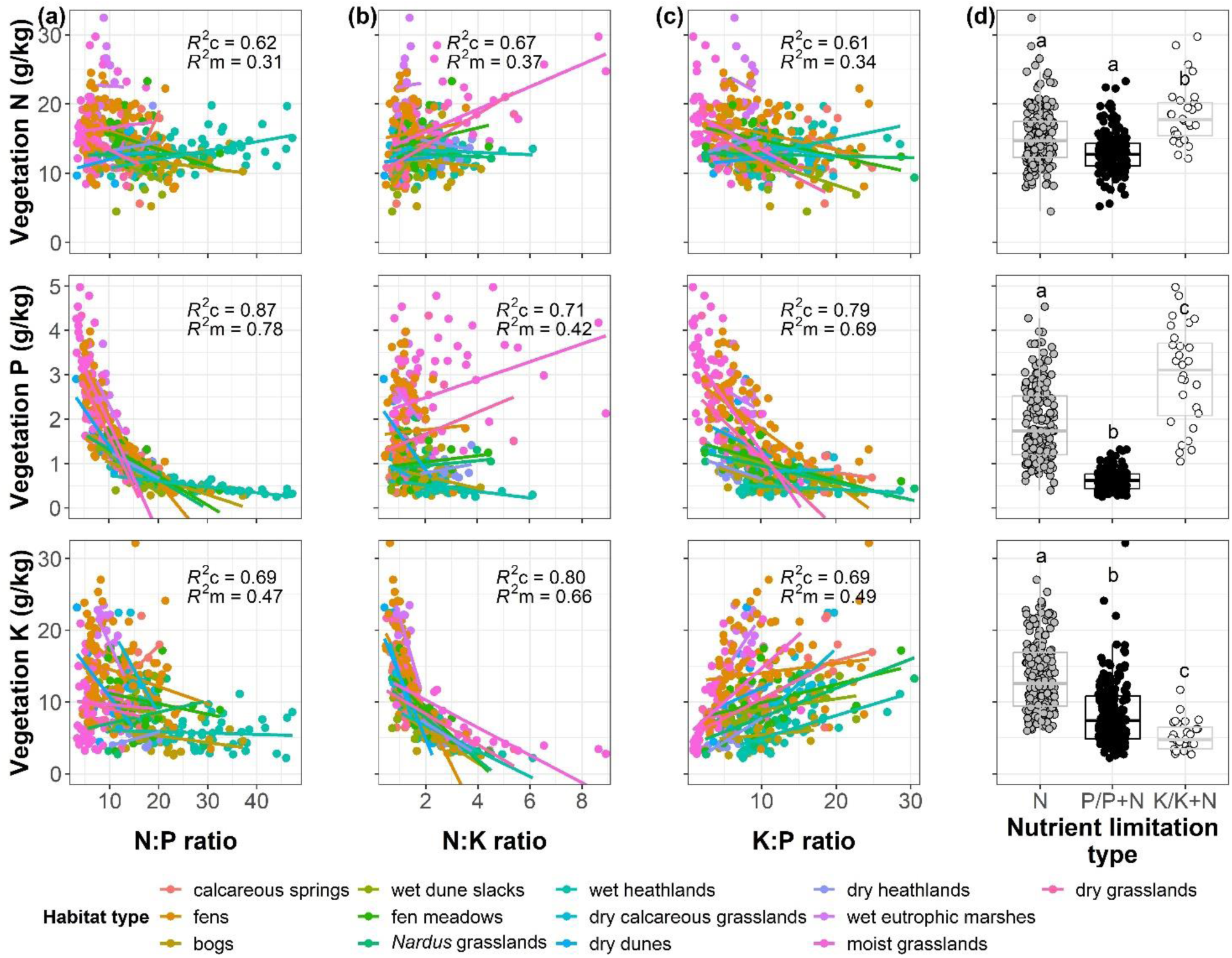
Variation in vegetation N:P:K stoichiometry (g g^−1^) and limitation types explained by concentrations of individual nutrients. a, the N:P ratio is correlated with P > K > N. b, the N:K ratio is correlated with K > P > N. c, the K:P ratio is correlated with P > K > N. d, P/P+N limited vegetation exhibits low P concentrations and K/K+N limited vegetation exhibits low K concentrations (in combination with high N and P), but N limited vegetation is not associated with low N concentrations.

Investigating correlations between vegetation concentrations and their ratios further reveals which nutrient was the strongest driver of the stoichiometry and its derived limitation types (Fig. 4a-c; Extended Data Table 4). Across the dataset (and while accounting for differences in relationships among habitat types), P clearly and strongly correlated with N:P, P:N, P:K and K:P ratios, more so than the other nutrients in the respective ratios. Regarding N:K and K:N, K correlated the strongest of the two nutrients, and P related even slightly better to N:K and K:N than N did. Taken together, this suggests that P affects the ratios stronger than K, and K stronger than N. Combined with evidence from the boxplots, this translates to P as the primary determinant of N versus P/P+N limitation, more so than N. As for K, most of the observed range of concentrations and ratios with K were not associated with K/K+N limitation. Instead, K/K+N limitation occurred where moderately low K concentrations are combined with both high N and high P.

## Discussion

Of the biodiverse herbaceous communities that we investigated, about as many are P (co)limited as there are limited by N (Figs. 1 and 2b). This finding supports the hypothesis that P (co)limitation in temperate regions is not restricted to aquatic ecosystems, and that it frequently occurs in terrestrial temperate ecosystems, too^6^. With respect to N and P, our results support that the Walker and Syers model of an evolution from N to P limitations with soil development^32,47^ remains valid over a variety of habitat types: the availability of N increases relative to that of P from earlier successional habitats in terms of soil age (soil total N = 0.2-1.0 g kg^−1^), such as dry dunes and dune slacks^48^, to later successional habitats in terms of soils (soil total N = 0.3-26 g kg^−1^; Table S1) that include *Nardus* grasslands, dry and wet heathlands^49,50^ (Table 1 and Fig. 2). Consequently, N limitation shifts toward P/P+N limitation along the soil development gradient. For soil pH, the expectation of optimal N and

P availabilities around slightly below neutral acidity holds, with optima around pH = 5.6 and pH = 6.0 for vegetation N and P, respectively (Table 1 and Fig. 3b). However, some noteworthy and unexpected patterns emerge in the availability and (co)limitation of K, the dominance of N versus P versus K as the driver of nutrient stoichiometries and limitations, and in the importance of soil moisture versus pH.

N:K ratios and K/K+N limitation decrease with increasing soil moisture as anticipated (Table 1), but a stronger trend in the same direction for N:K-soil pH and K/K+N-soil pH associations, as well as a positive K:P-soil pH relationship above pH = 4.3 (Extended Data Fig. 3) suggests that not only an inhibited N mineralization at high soil moisture^51^ is at play here. Instead, combining the patterns along the moisture and acidity axes points to the increase of K availability due to input of base-rich (ground)water. Such conditions occur in calcareous springs, (alkaline) fens and wet eutrophic marshes (Extended Data Fig. 1). The association with pH here looks stronger than that with soil moisture, because irrespective of average moisture, soil pH controls the retention of base cations such as K on cation exchange sites^52^, and the other way around, base cations buffer soil pH.

Nevertheless, also at the lower end of the pH spectrum, where habitats are found with minimal vegetation K concentration (e.g. bogs, dry and wet heathlands - Extended Data Fig. 1b and Fig. S1), actual (co)limitation by K remains rare in biodiverse communities without historical heavy fertilization, as studied here (Figs. 1c and 2b). This was also the case in earlier studies that observed K (co)limitation only sporadically in temperate wetlands and other herbaceous habitats^13,18,53^. Only where the vegetation becomes species-poorer under higher P (and N) loads (moist and dry grasslands in our study), K/K+N limitation is a more common occurrence (Fig. 2b). K/K+N limitation thus seems to emerge where the availability of P (and to a lesser extent N) is high, making K simply the next limiting nutrient in line^54^. Supporting this interpretation, we found a positive effect of N deposition on vegetation N:K, but this only resulted in K/K+N limitation when testing specifically among the already more species-impoverished or eutrophic habitats (Table 1). Related, K/K+N limitation occurs only where both vegetation K is relatively (but not minimally) low and at the same time vegetation contents of N and particularly P are high (Fig. 4). Our results on K/K+N limitation do therefore not necessarily disagree with reports on widespread K (co)limitation, based on K fertilization experiments: such meta-analyses^52,55^ or syntheses of coordinated distributed experiments^8^ usually do not exclude already (formerly) nutrient-enriched habitats degraded in species richness *a priori*.

P/P+N limitation appears where P availability is at its lowest, as opposed to N limited communities which do not differ in their N contents from P/P+N limited vegetation. Correlating the vegetation nutrient concentrations with their ratios (Fig. 4) then reveals that not the availability of N, but that of P is the strongest determinant of the N:P ratio. Similarly, vegetation P is more strongly correlated with P:K ratios than is K, but K is still the stronger determinant of the N:K ratio in comparison to N. Stronger plant P versus N:P than N versus N:P correlations have earlier been reported for within-species patterns across sites as well as within-site patterns across species^5^, but to our knowledge not for N, P and K at the vegetation level across biodiverse sites, spanning long gradients in soil development stages, moisture and pH. Güsewell (2004)^5^ ascribed the within-species differences in correlations to the relative stability of graminoid N, as opposed to woody plants, mosses and lichens for which N would explain most of N:P. At the sites we sampled, there is sevenfold variation in N concentrations at the vegetation level (range 4.5-32.5 g kg^−1^), compared to twentyfold variation in P (0.25-5 g kg^−1^) and fifteenfold variation in K (2.2-32 g kg^−1^). Hence, variation in vegetation N is effectively lowest, but still substantial.

The importance of P > K > N in determining vegetation stoichiometry translates further into the influence of the environmental pH gradient. Soil pH has a non-linear influence on both N and P availability (and a near-linear, positive effect on K), with optima around pH = 5.6 and 5.8 respectively. With the pH influences on the two nutrients combined, the N:P ratio is minimal at around pH = 6.4 (Fig. 3a). Both the convex nature of the curve and the position of the minimum indicate that the physicochemistry-driven^56^ positive effect of near-neutral pH on P is more important than the microbially-driven^27^ positive effect of slightly below neutral pH on N. Also maximum N limitation at pH = 6.1 and minimum P/P+N limitation at pH = 6.0 (Fig. 3b) point to an effect of pH on vegetation nutrient limitation primarily through affecting P availability, rather than N. The greater importance of optimal pH to P than to N availability implies that in high biodiversity herbaceous vegetation without heavy or long-term fertilization, P is (co)limiting at low and high pH, whereas N is limiting at intermediate acidity.

Relationships between soil moisture and nutrient concentrations, ratios and limitation types were weaker than anticipated, in particular when compared to these with soil pH (Table 1). For instance, the hypothesized reduced decomposition and nitrification rates with increasing moisture^25,51^ did not result in decreased vegetation N. Similarly, the hypothesized positive effect of moisture on vegetation P and K through improved molecular diffusion^42^ was absent for P, and only weak and of minor importance for K. Related, P/P+N limitation occurs even slightly more commonly at high than at low soil moisture, opposed to what we expected. A contributing aspect here may be that at some of the dry sites (heathlands, dunes, calcareous and non-calcareous grasslands – Extended Data Fig. 1a and Extended Data Table 1) but never at the wet sites, water might in reality be a more strongly limiting resource than nutrients^57^. Consequently, biodiverse non-light limited vegetation can develop here without strong nutrient constraints. Vegetation growing in moister conditions (excluding waterlogged situations with anoxia and toxicity stress^1^), on the other hand, will only be biodiverse and non-light limited if effectively nutrients are limiting. In practice this will often be limitation by P/P+N. Nonetheless, for the habitats at both the wetter and drier ends of the moisture spectrum, a mixture of nutrient limitation types was found (Fig. 2b), and on the drier side, while water limitation can occur, it is known from fertilization experiments that many heathlands are effectively limited by P/P+N^58^, while decalcified dune vegetation^59^ and temperate grasslands^60^ are limited by N.

Soil moisture further interacts negatively with pH in influencing vegetation N (Extended Data Table 2), indicating that the influence of pH on N, showing an optimum, becomes more important with increasing soil moisture. However, also here, when testing for effects of both soil moisture and pH together (and potentially interacting), variation explained by soil moisture is low compared to that by pH. The fact that soil pH, with its clearer influence on P than on N, also relates much more strongly to stoichiometric ratios and limitation types than moisture, indicates again the primary role of P availability and its controls in influencing nutrient limitation.

Across our dataset spanning strong contrasts in habitats in terms of environmental conditions and community composition, spatial variation in atmospheric N deposition played a secondary role in modifying vegetation stoichiometry and limitation. When testing for the influence of N deposition in isolation, log N:K is found to increase on average by 0.03 units per additional kg of N load ha^−1^yr^−1^ (Table 1). Accordingly, N deposition promotes the occurrence of K/K+N limitation, especially in the already more eutrophic or species-impoverished habitats (dry and moist grasslands – Figs. 1a and 2b, Table 1). Such N deposition-only effect was not found on the N:P ratio nor on the occurrence of N vs P/P+N limitation (Table 1), but when estimating the combined influence of soil moisture and measured pH on N:P, simultaneously accounting for N deposition does substantially improve the variation in N:P explained (from *R²*_m_ = 0.22 to *R²*_m_ = 0.32 – Extended Data Tables 2 vs 3). An interactive effect between soil pH and N deposition is suggested here, with the influence of the N load on N:P becoming more important under acidic conditions (e.g. an N:P increase by 0.5 units per additional kg N ha^−1^yr^−1^ at pH = 4, and N:P increase by 0.2 per additional kg N ha^−1^yr^−1^ at pH = 6), potentially further reinforcing P/P+N limitation particularly at low pH. This result aligns with findings by Aerts et al. (1992)^28^, who observed significantly increased N:P ratios with natural and manipulated N deposition in acidic bogs, and a meta-analysis of N addition experiments by Xu et al. (2022)^61^ who found more pronounced N:P increases where soil buffer capacity is lowest.

In conclusion, soil pH plays a dominant role in determining stoichiometry and limitation types of biodiverse, herbaceous plant communities. External and internal eutrophication further significantly and meaningfully modifies these vegetation N:P:K ratios; where single or combined impacts of eutrophication sources such as N deposition and land use legacies (e.g. in old fields^8^) are of such extent that also plant biodiversity is clearly affected, naturally rarer types of nutrient limitation, such as by K/K+N, appear more often. Finally, observed distinctions in nutrient limitation types among habitats underscore the necessity to consider the environmental conditions of each ecosystem, both from a fundamental scientific point of view (e.g. caution with extrapolations of results between less and more disturbed systems, and between those with optimal versus suboptimal soil pH) as well as from an applied perspective, requiring habitat-specific management approaches.

## Methods

### Data collection and processing

We sampled 386 low productive herbaceous ecosystems across temperate northwest and central Europe (Table S1), for which nutrients are the most likely factor limiting growth and determining vegetation processes (e.g. standing crop was < 6000 kg ha^−1^ for 87% of sites, suggesting minor prevalence of light limitation^44^). 287 of the sites harbored biodiverse herbaceous vegetation corresponding to diverse Natura 2000 habitat types, and 99 others exhibited less species rich communities or communities that classified not straightforwardly into such habitat type (Fig. S2). At all sites, we determined herbaceous community aboveground N, P and K concentrations at the peak of the growing season, prior to grazing or mowing (if any). General soil moisture conditions were expressed as the Ellenberg Indicator Value (EIV) for soil moisture^62,63^, which had been estimated for all sites based on species presence-absence data (and not abundance because of variation in the Braun-Blanquet, Londo and percentual methods to estimate cover during data collection^64^) in corresponding relevees. The EIV for soil pH was determined analogously to that of moisture, but in addition, pH was measured for 327 of the sites - in the field or lab, in water or in KCl-solution (correlation between EIV and measured pH: *r* = 0.87). Intra-site spatial replicates of nutrient data were averaged, and for inter-annual temporal replicates, we used the most recent data point.

#### Type of nutrient limitation

At each site in summer (June-August), aboveground parts of all vascular plants within a 40×40 cm square were sampled to determine mass-based nutrient concentrations. Litter was excluded, but tissue senesced during the growing season of sampling was included. Collected plant material was dried for at least 72 hours at 70°C and weighed. Subsamples were subsequently ground for analysis of N, P and K (g kg^−1^ plant material) through acid digestion^65^ with sulfuric acid, salicylic acid, hydrogen peroxide and selenium, and measured with a segmented flow analyzer (SAN++, Skalar, Breda, NL).

Based on the mass-based elemental ratios and the critical values as defined by Olde Venterink et al. (2003)^13^, we categorized the samples into following types of nutrient limitation:

limitation: N:P < 14.5 and N:K < 2.1
P- or P+N limitation: N:P > 14.5 and K:P > 3.4
K- or K+N limitation: N:K > 2.1 and K:P < 3.4

For the large majority of sites (96%) we were able to determine the type of nutrient limitation in this way.

#### Soil moisture

Overall soil moisture regimes were determined indirectly based on the presence and absence of plant species within larger sampling plots that varied in size between 4 and 25 m² (Table S2). With help of the JUICE software^66^ in combination with EIVs from Berg (2011)^67^ for Polish sites (now also available at Europe-wide scale^63^), and Turboveg software^68^ in combination with EIVs from Ellenberg et al. (1992)^62^ for the other sites, soil moisture regimes were assigned values between 3 (most dry) and 10 (most wet). We considered plots and habitats with a moisture EIV < 5.5 as dry, with an EIV of 5.5–8.0 as moist, and with an EIV > 8.0 as wet.

#### Soil pH and total nitrogen

Analogous to the methodology for soil moisture, the acidity of the soil was also estimated indirectly based on species’ presence and their corresponding EIVs for pH, ranging between 1 (most acidic) and 9 (most alkaline). In addition, for 327 of the sites, pH of the upper 5 cm of soil was determined either in the field with a pH meter (Hanna Instruments), or afterwards with a soil sample in water (pH H_2_O) or KCl solution (pH KCl)^69^. Since pH H_2_O and pH KCl are not interchangeable, and we aimed to take both types of measurements as well as the EIV into account in some analyses for data coverage purposes, we decided to work with a pH classification. To this end, we based ourselves on the classification system used for Habitat 2000 areas by the Dutch Ministry of Agriculture, Nature and Food Quality^70^. In cases where both pH KCl and pH H_2_O were determined but disagreed on the categorization, we opted for pH KCl. Furthermore, being direct estimates, measured pH values were always preferred over EIVs where available:

pH class 1 (acidic): pH KCl < 3.5, pH H_2_O < 4.5, EIV pH ≤ 3
pH class 2 (moderately acidic): pH KCl 3.5–4.8, pH H_2_O 4.5–5.5, EIV pH 3–4.5
pH class 3 (weakly acidic): pH KCl 4.8–6.1, pH H_2_O 5.5–6.5, EIV
pH 4.5–5.9 pH class 4 (circumneutral): pH KCl 6.1–7.5, pH H_2_O 6.5–7.5, EIV
pH 5.9–7.3 pH class 5 (alkaline): pH KCl > 7.5, pH H_2_O > 7.5, EIV pH > 7.3

Besides the pH, also total N concentrations of 0–5 cm soil samples were determined for 230 of the sites (Table S1). Soil total N was used in the present study as further supporting evidence (in combination with knowledge based on expertise and literature) to classify habitat types in terms of their soils as young or further developed. These N contents were determined with an elemental analyzer (Flash EA 2000, Thermo Fisher Scientific, Bremen, DE) or through acid digestion^65^ with a segmented flow analyzer (SAN++, Skalar, Breda, NL).

*Habitat types*. Based on the environmental gradients in soil moisture and pH, we established a raster with the 13 habitat types investigated in the present study (Extended Data Table 1). These habitat types, further detailed in Table S3, were mostly assigned in-situ based on species’ presence and ecology^71^. For plots not classified locally, we determined the habitat type based on the location and the presence of vegetation characteristic species, following ref.^72^. To this end, we performed a redundance analysis (RDA) with help of the R ‘vegan’ package^73^. Characteristic species of the *Ericetum tetralicis*, *Calluno-Geniston pilosae*, *Junco-Molinion*, *Nardo-Galion Saxatilis*, *Caricetalia nigrae*, *Thero-Airion* and *Erico-Sphagnetum magellanici* were used to distinguish, respectively, wet heathlands, dry heathlands, *Molinia* meadows (of category fen meadows), *Nardus* grasslands, wet dune slacks, inland dunes (of category dry dunes) or raised bogs (of category bogs).

In some plots, the habitat was clearly in a degraded and species-impoverished state, or a classification into one of the first ten Natura 2000 habitat types was not possible. Such more eutrophic or eutrophied habitats were classified as ‘wet eutrophic marshes’ (i.e. eutrophic marshes with *Filipendula ulmaria* or reedbeds), ‘moist grasslands’ (i.e. moist or wet grasslands or meadows not in Natura 2000), or ‘dry grasslands’ (i.e. dry grasslands or meadows not in Natura 2000).

#### Nitrogen deposition

N deposition for each site was determined based on high resolution (2 x 2 km² in northwest Europe (2-16°E, 47-56°N), 25 x 25 km² elsewhere in Europe) estimates applicable to its coordinates (Fig. S3). Specifically, we summed annual dry and wet deposition modelled with the LOTOS-EUROS model v2.2.002^74^ for the readily available values of the years 2000, 2005, 2010, 2015 and 2019. The model simulations setup used 12 vertical levels, extending from the ground to about 10 km altitude. The emissions used at the coarser European scale were based on the CAMS-REGv5.1 inventory^75^ for all non-2000 years, and the CAMSv5.1 inventory for 2000. The higher resolution model run used a combination of CAMS-REGv5.1 and the more detailed GrETa inventory^76^. For dry deposition the DEPAC model was used^77^, which includes a compensation point approach for NH_3_ ^78^. Wet deposition was parameterized in terms of size-dependent scavenging and collection efficiencies for particles and corresponding uptake coefficients for gases^74^. The deposition of N in HNO_3_, NO, NO_2_, NH_3_, aerosol phase NO ^−^ and aerosol phase NH ^+^ were summed for each location and subtracted by dry NH_3_ emission. For each site, the N deposition estimate for the last available year prior to sampling was used, except for eight data points from 1999 that were assigned 2000 estimates, avoiding the use of lower-resolution pre-2000 data.

### Statistical analysis

We plotted ternary diagrams with help of the R package ‘ggtern’^79^ to visualize, explore and describe our data. Color codes for data points in the diagrams distinguished habitat types or values for the ecological gradients in soil moisture, pH and N deposition.

Linear mixed modeling in R version 4.3.3^80^ with package lmerTest^81^ was used to formally analyze our research questions on variations in nutrient concentrations and stoichiometry among habitat types, soil development stages and ecological gradients. Also differences in soil moisture EIVs and pH classes among habitat types were tested in this way (taking into account the ordinal nature of pH classes with the clmm function of the R package ordinal^82^). The areas (locations) in which individual data points (sites) were located, were accounted for in the random effects (e.g. ‘Torfbroek_fens’, ‘Torfbroek_fenmeadows’, ‘Biebrza_fens’, ‘Liereman_wet_heath’): if the distance between two consecutive sites was > 3000 m, or the habitat type differed, these were considered independent sites, i.e. they were assigned a different location name. This way, we distinguished 142 locations to control for variables not taken into account in the analysis, such as geological and historical backgrounds. This procedure effectively resulted in mixed model residuals that did not exhibit spatial autocorrelation according to Moran’s I test (moran.test from R package spdep^83^).

In order to analyze effects of environmental gradients on nutrient limitation types, we adopted a generalized linear mixed modeling approach with binomial distribution of the response variable. For this, the nutrient limitation type of interest (N, P/P+N or K/K+N) was coded as 1, with the other two limitation types coded as 0. This generalized modeling approach was also followed when testing for differences in limitation types among habitat types, but here, in combination with observing visual variations in bar charts, a fixed model without the random ‘location’ term was used as an approximation because of convergence issues with the mixed model approach.

For multiple regressions, testing the combined effects of soil moisture, pH and N deposition, the model selection procedure started with a full model with all factors (including a potential quadratic pH effect), and two-way interactions if suggested by visual inspection of regression trees (functions tree and prune.tree of the tree package in R^84^). Next, models were simplified by removing least-significant terms one by one, until only significant predictors remained.

N, P and K concentrations were set out against their ratios and the limitation types to unravel whether stoichiometry and limitations are predominantly influenced by the availability of either nutrient. To this end, nutrient concentrations were plotted against ratios, while accounting for varying slopes among habitat types (fixed) as well as shared locations (random). Marginal and conditional *R²* here provide an idea of to what extent the ratios are driven by the different nutrients. For the nutrient limitations, boxplots visualize the range of concentrations by limitation type. Differences here among the three limitation types were tested analogously to the mixed modeling approach explained above.

Linear model assumptions were checked through histograms, the Shapiro-Wilk normality test on residual values, and plots of these residuals versus fitted values. For generalized linear models, assumptions of uniformity, absence of outliers, overdispersion and zero inflation and testing of residual versus fitted values were performed with tests provided in the DHARMa R package^85^. Where necessary to meet assumptions, the response variable was transformed (e.g. N:K was log-transformed to address a positively skewed distribution). The significance level was α = 0.05, and *P* < 0.10 was interpreted as near-significant. Marginal and conditional (partial) proportions of variation explained by mixed models were estimated with the partR2 function^86^. The R function lsmeans with Tukey’s post hoc correction for multiple testing was used for detecting and quantifying significant between-group differences^87^.

## Data availability

Data are presented in Tables S1-S3 as well as on: github.com/KevinVanSundert/NPK_NatPlants_2024_KVS

## Code availability

The code and data to reproduce all results are available on: github.com/KevinVanSundert/NPK_NatPlants_2024_KVS

## Acknowledgements

This research was supported by BiodivERsA project REPEAT. KVS and WJE acknowledge support from the Research Foundation-Flanders (FWO awards no. 1222323N and 1214520N).

## Extended data

### Characterization of habitat types

**Extended Data Fig. 1.**
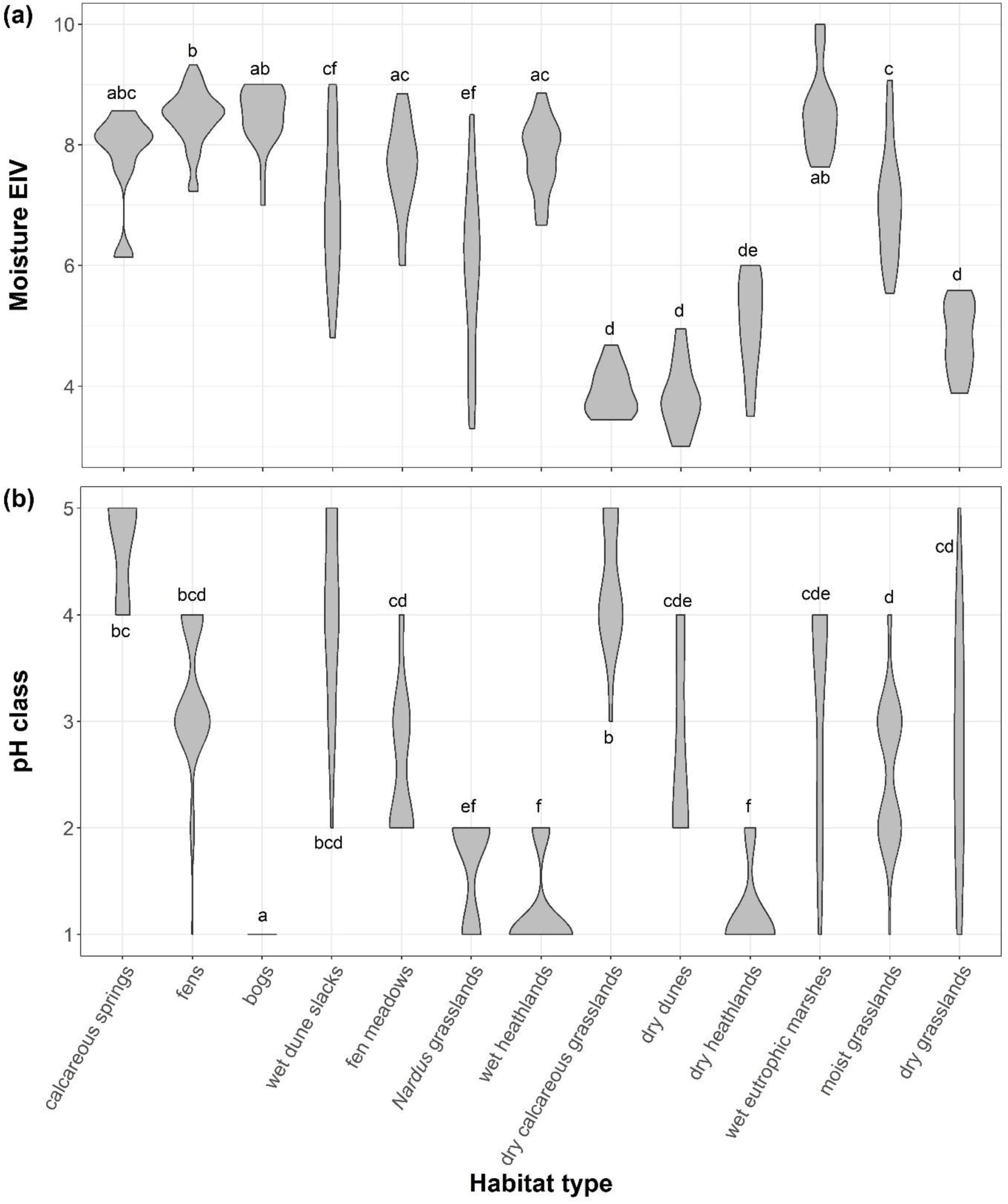
Distribution of environmental gradients per habitat type. **a**, Ellenberg Indicator Values (EIV) for soil moisture. **b**, pH classes where 1 = acidic, 2 = moderately acidic, 3 = weakly acidic, 4 = circumneutral, 5 = alkaline.

**Extended Data Table 1.** Habitat types classified along environmental gradients in soil moisture and pH.

| Soil pH / moisture | Wet | Moist | Dry |
| --- | --- | --- | --- |
| Alkaline/circumneutral | calcareous springs<br>wet eutrophic marshes | wet duneslacks | dry calcareous grasslands |
| Weakly/<br>moderately acidic | fens | fen meadows<br>*Nardus* grasslands<br>moist grasslands | dry dunes<br>dry grasslands |
| Acidic | bogs | wet heathlands | dry heathlands |

### Nutrient stoichiometry and limitation among habitat types

**Extended Data Fig. 2.**
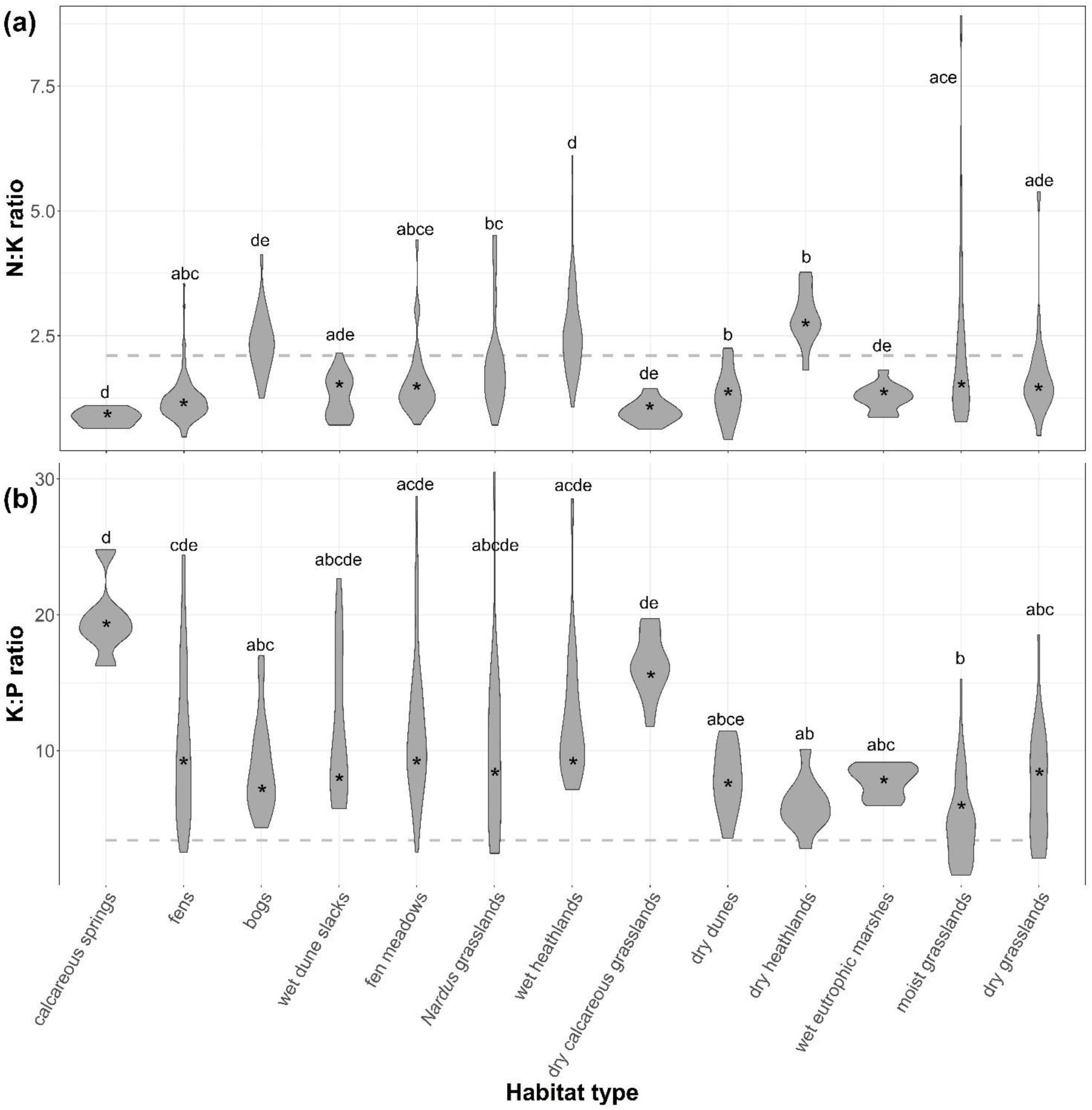
Patterns of vegetation N:K and K:P stoichiometry (g g^−1^) across habitat types. **a**, the N:K ratio, with critical ratio of 2.1^13^ indicated through a dashed line. **b**, the K:P ratio, with critical ratio of 3.4 indicated through a dashed line. Values significantly differing from the respective critical ratios were marked with ‘*’, values significantly different among habitat types were indicated trough different letters.

### Nutrient limitation and environmental gradients

**Extended Data Fig. 3.**
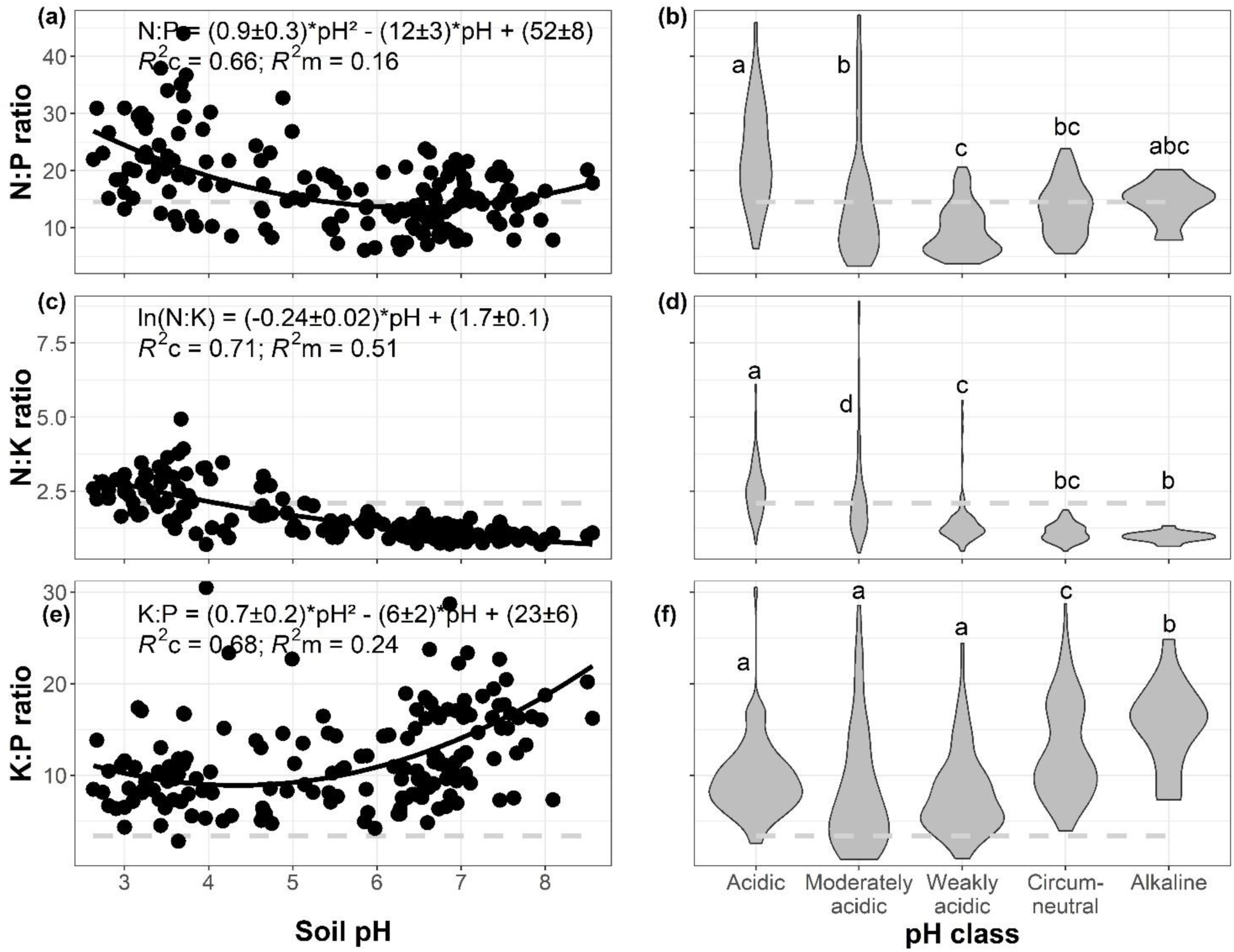
Associations of vegetation N:P:K stoichiometry (g g^−1^) with soil pH. **a**, **c**, **e**, ratios set out against soil pH measured in water. **b**, **d**, **f**, ratios set out against classified pH. Critical ratios of N:P (14.5), N:K (2.1) and K:P (3.4)^13^ are indicated through dashed lines.

**Extended Data Table 2.**
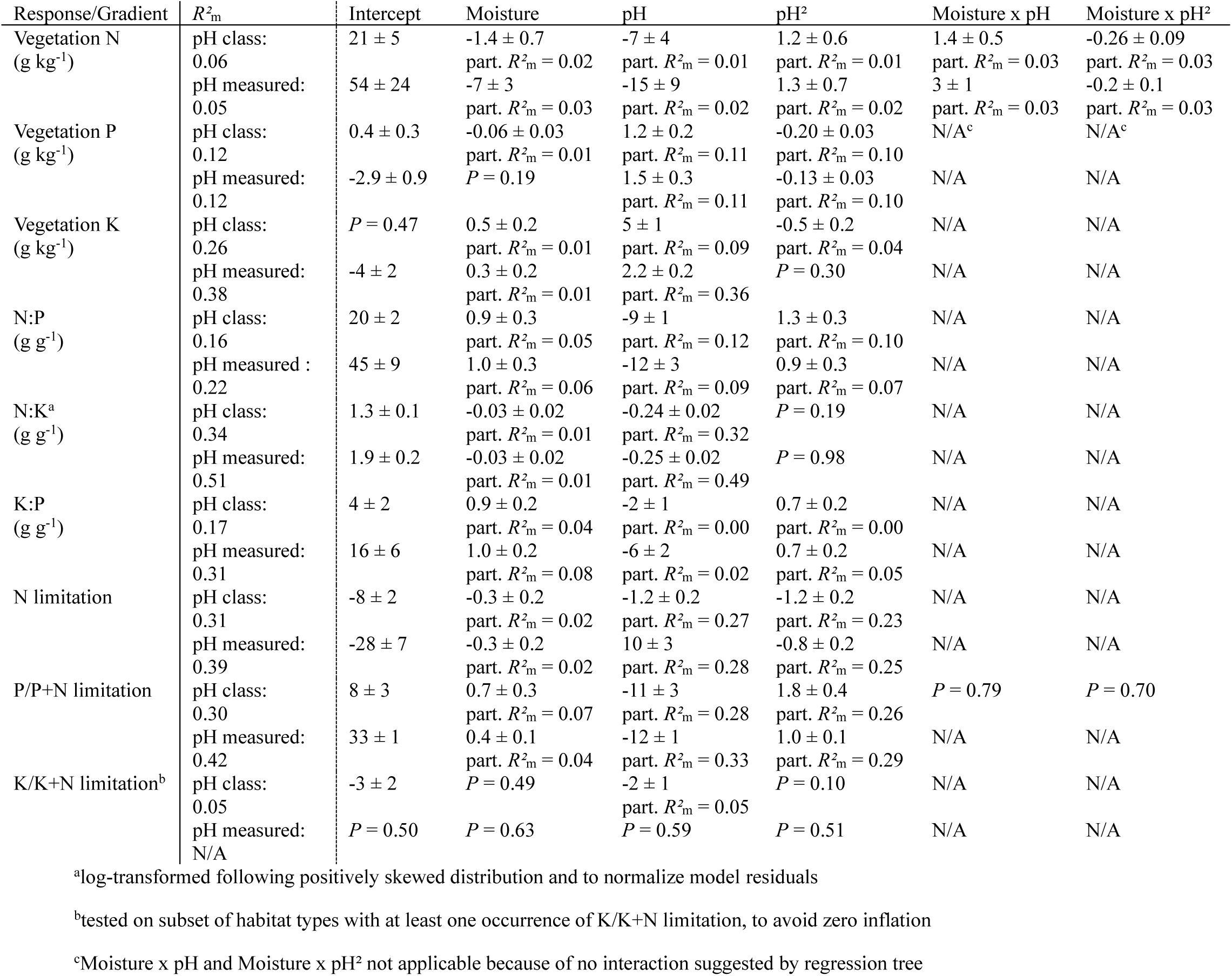
Associations of vegetation N, P and K concentrations, stoichiometry and limitation types with combined environmental gradients in soil moisture and pH.

**Extended Data Table 3.**
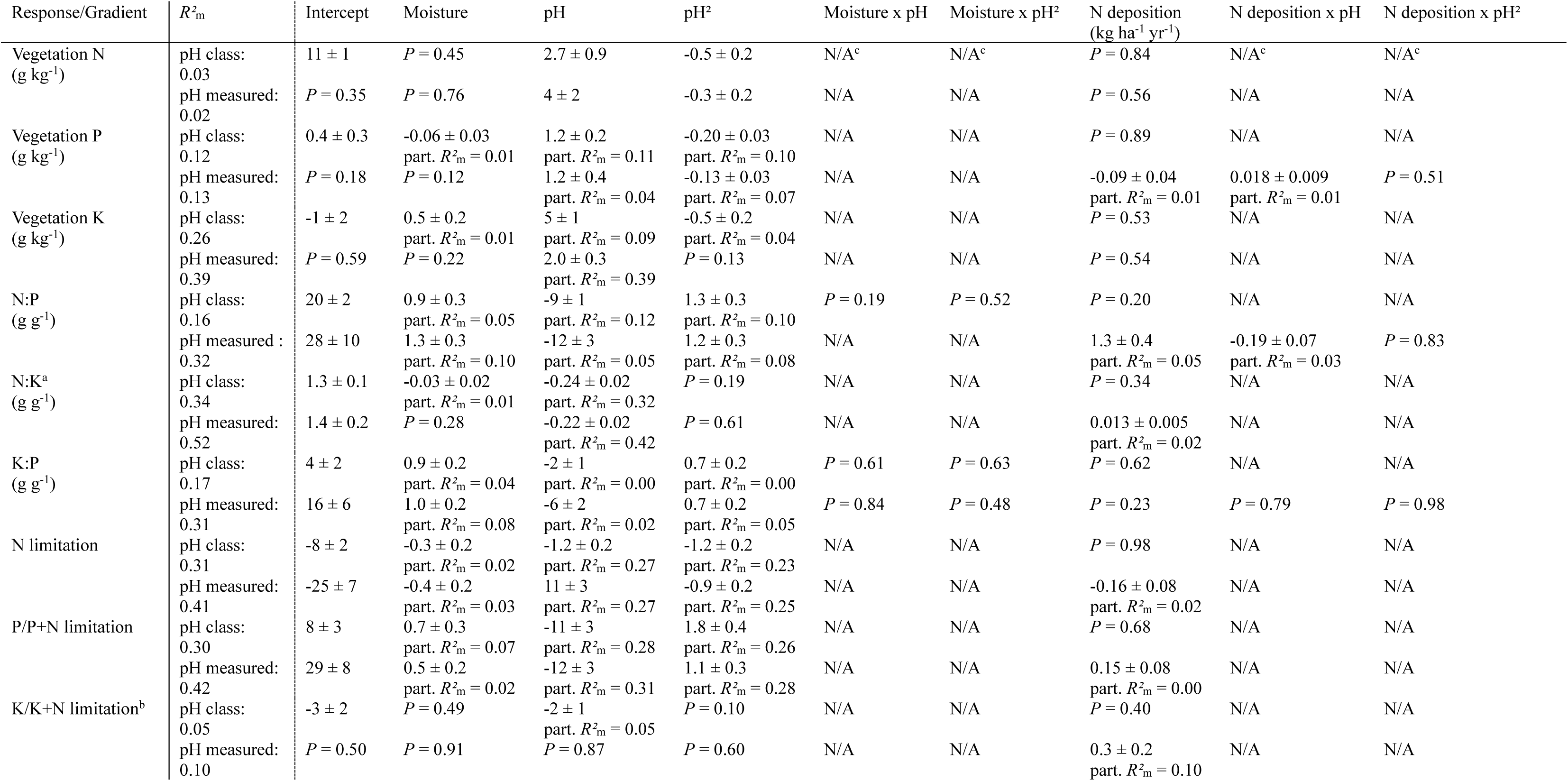
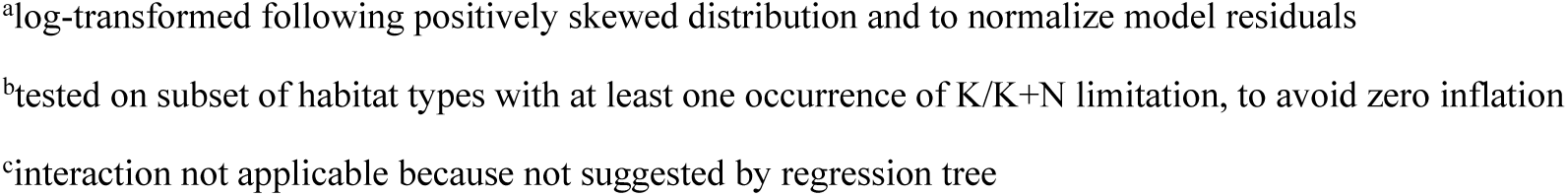
Associations of vegetation N, P and K concentrations, stoichiometry and limitation types with combined environmental gradients in soil moisture and pH, while accounting for N deposition effects.

| Response/Gradient | $R^2_m$ | Intercept | Moisture | pH | pH<sup>2</sup> | Moisture x pH | Moisture x pH<sup>2</sup> | N deposition<br>(kg ha<sup>-1</sup> yr<sup>-1</sup>) | N deposition x pH | N deposition x pH<sup>2</sup> |
| --- | --- | --- | --- | --- | --- | --- | --- | --- | --- | --- |
| Vegetation N<br>(g kg<sup>-1</sup>) | pH class:<br>0.03 | 11 ± 1 | $P = 0.45$ | 2.7 ± 0.9 | -0.5 ± 0.2 | N/A<sup>c</sup> | N/A<sup>c</sup> | $P = 0.84$ | N/A<sup>c</sup> | N/A<sup>c</sup> |
| | pH measured:<br>0.02 | $P = 0.35$ | $P = 0.76$ | 4 ± 2 | -0.3 ± 0.2 | N/A | N/A | $P = 0.56$ | N/A | N/A |
| Vegetation P<br>(g kg<sup>-1</sup>) | pH class:<br>0.12 | 0.4 ± 0.3 | -0.06 ± 0.03<br>part. $R^2_m = 0.01$ | 1.2 ± 0.2<br>part. $R^2_m = 0.11$ | -0.20 ± 0.03<br>part. $R^2_m = 0.10$ | N/A | N/A | $P = 0.89$ | N/A | N/A |
| | pH measured:<br>0.13 | $P = 0.18$ | $P = 0.12$ | 1.2 ± 0.4<br>part. $R^2_m = 0.04$ | -0.13 ± 0.03<br>part. $R^2_m = 0.07$ | N/A | N/A | -0.09 ± 0.04<br>part. $R^2_m = 0.01$ | 0.018 ± 0.009<br>part. $R^2_m = 0.01$ | $P = 0.51$ |
| Vegetation K<br>(g kg<sup>-1</sup>) | pH class:<br>0.26 | -1 ± 2 | 0.5 ± 0.2<br>part. $R^2_m = 0.01$ | 5 ± 1<br>part. $R^2_m = 0.09$ | -0.5 ± 0.2<br>part. $R^2_m = 0.04$ | N/A | N/A | $P = 0.53$ | N/A | N/A |
| | pH measured:<br>0.39 | $P = 0.59$ | $P = 0.22$ | 2.0 ± 0.3<br>part. $R^2_m = 0.39$ | $P = 0.13$ | N/A | N/A | $P = 0.54$ | N/A | N/A |
| N:P<br>(g g<sup>-1</sup>) | pH class:<br>0.16 | 20 ± 2 | 0.9 ± 0.3<br>part. $R^2_m = 0.05$ | -9 ± 1<br>part. $R^2_m = 0.12$ | 1.3 ± 0.3<br>part. $R^2_m = 0.10$ | $P = 0.19$ | $P = 0.52$ | $P = 0.20$ | N/A | N/A |
| | pH measured :<br>0.32 | 28 ± 10 | 1.3 ± 0.3<br>part. $R^2_m = 0.10$ | -12 ± 3<br>part. $R^2_m = 0.05$ | 1.2 ± 0.3<br>part. $R^2_m = 0.08$ | N/A | N/A | 1.3 ± 0.4<br>part. $R^2_m = 0.05$ | -0.19 ± 0.07<br>part. $R^2_m = 0.03$ | $P = 0.83$ |
| N:K<sup>a</sup><br>(g g<sup>-1</sup>) | pH class:<br>0.34 | 1.3 ± 0.1 | -0.03 ± 0.02<br>part. $R^2_m = 0.01$ | -0.24 ± 0.02<br>part. $R^2_m = 0.32$ | $P = 0.19$ | N/A | N/A | $P = 0.34$ | N/A | N/A |
| | pH measured:<br>0.52 | 1.4 ± 0.2 | $P = 0.28$ | -0.22 ± 0.02<br>part. $R^2_m = 0.42$ | $P = 0.61$ | N/A | N/A | 0.013 ± 0.005<br>part. $R^2_m = 0.02$ | N/A | N/A |
| K:P<br>(g g<sup>-1</sup>) | pH class:<br>0.17 | 4 ± 2 | 0.9 ± 0.2<br>part. $R^2_m = 0.04$ | -2 ± 1<br>part. $R^2_m = 0.00$ | 0.7 ± 0.2<br>part. $R^2_m = 0.00$ | $P = 0.61$ | $P = 0.63$ | $P = 0.62$ | N/A | N/A |
| | pH measured:<br>0.31 | 16 ± 6 | 1.0 ± 0.2<br>part. $R^2_m = 0.08$ | -6 ± 2<br>part. $R^2_m = 0.02$ | 0.7 ± 0.2<br>part. $R^2_m = 0.05$ | $P = 0.84$ | $P = 0.48$ | $P = 0.23$ | $P = 0.79$ | $P = 0.98$ |
| N limitation | pH class:<br>0.31 | -8 ± 2 | -0.3 ± 0.2<br>part. $R^2_m = 0.02$ | -1.2 ± 0.2<br>part. $R^2_m = 0.27$ | -1.2 ± 0.2<br>part. $R^2_m = 0.23$ | N/A | N/A | $P = 0.98$ | N/A | N/A |
| | pH measured:<br>0.41 | -25 ± 7 | -0.4 ± 0.2<br>part. $R^2_m = 0.03$ | 11 ± 3<br>part. $R^2_m = 0.27$ | -0.9 ± 0.2<br>part. $R^2_m = 0.25$ | N/A | N/A | -0.16 ± 0.08<br>part. $R^2_m = 0.02$ | N/A | N/A |
| P/P+N limitation | pH class:<br>0.30 | 8 ± 3 | 0.7 ± 0.3<br>part. $R^2_m = 0.07$ | -11 ± 3<br>part. $R^2_m = 0.28$ | 1.8 ± 0.4<br>part. $R^2_m = 0.26$ | N/A | N/A | $P = 0.68$ | N/A | N/A |
| | pH measured:<br>0.42 | 29 ± 8 | 0.5 ± 0.2<br>part. $R^2_m = 0.02$ | -12 ± 3<br>part. $R^2_m = 0.31$ | 1.1 ± 0.3<br>part. $R^2_m = 0.28$ | N/A | N/A | 0.15 ± 0.08<br>part. $R^2_m = 0.00$ | N/A | N/A |
| K/K+N limitation<sup>b</sup> | pH class:<br>0.05 | -3 ± 2 | $P = 0.49$ | -2 ± 1<br>part. $R^2_m = 0.05$ | $P = 0.10$ | N/A | N/A | $P = 0.40$ | N/A | N/A |
| | pH measured:<br>0.10 | $P = 0.50$ | $P = 0.91$ | $P = 0.87$ | $P = 0.60$ | N/A | N/A | 0.3 ± 0.2<br>part. $R^2_m = 0.10$ | N/A | N/A |

<sup>a</sup>log-transformed following positively skewed distribution and to normalize model residuals
<sup>b</sup>tested on subset of habitat types with at least one occurrence of K/K+N limitation, to avoid zero inflation
<sup>c</sup>interaction not applicable because not suggested by regression tree

### Which nutrient drives the limitation?

**Extended Data Table 4.** Variation in vegetation N:P:K stoichiometry (g g^−1^) explained by concentrations of individual nutrients.

| Ratio/Concentration | N ($\text{g kg}^{-1}$) | P ($\text{g kg}^{-1}$) | K ($\text{g kg}^{-1}$) |
| --- | --- | --- | --- |
| N:P | $R^2_{\text{m}} = 0.31$ | $R^2_{\text{m}} = 0.78$ | $R^2_{\text{m}} = 0.47$ |
| | $R^2_{\text{c}} = 0.62$ | $R^2_{\text{c}} = 0.87$ | $R^2_{\text{c}} = 0.69$ |
| P:N | $R^2_{\text{m}} = 0.29$ | $R^2_{\text{m}} = 0.83$ | $R^2_{\text{m}} = 0.47$ |
| | $R^2_{\text{c}} = 0.61$ | $R^2_{\text{c}} = 0.90$ | $R^2_{\text{c}} = 0.68$ |
| P:K | $R^2_{\text{m}} = 0.36$ | $R^2_{\text{m}} = 0.68$ | $R^2_{\text{m}} = 0.51$ |
| | $R^2_{\text{c}} = 0.62$ | $R^2_{\text{c}} = 0.77$ | $R^2_{\text{c}} = 0.71$ |
| K:P | $R^2_{\text{m}} = 0.34$ | $R^2_{\text{m}} = 0.69$ | $R^2_{\text{m}} = 0.49$ |
| | $R^2_{\text{c}} = 0.61$ | $R^2_{\text{c}} = 0.79$ | $R^2_{\text{c}} = 0.69$ |
| N:K | $R^2_{\text{m}} = 0.37$ | $R^2_{\text{m}} = 0.42$ | $R^2_{\text{m}} = 0.66$ |
| | $R^2_{\text{c}} = 0.67$ | $R^2_{\text{c}} = 0.71$ | $R^2_{\text{c}} = 0.80$ |
| K:N | $R^2_{\text{m}} = 0.34$ | $R^2_{\text{m}} = 0.44$ | $R^2_{\text{m}} = 0.74$ |
| | $R^2_{\text{c}} = 0.62$ | $R^2_{\text{c}} = 0.72$ | $R^2_{\text{c}} = 0.86$ |

## References

1. Kotowski, W., Beauchard, O., Opdekamp, W., Meire, P. & Van Diggelen, R. Waterlogging and canopy interact to control species recruitment in floodplains. Funct. Ecol. 24, 918–926 (2010).

2. Grime, J. P. Plant Strategies and Vegetation Processes. (Wiley, 1979).

3. Vitousek, P. & Howarth, R. Nitrogen limitation on land and in the sea: How can it occur? Biogeochemistry 13, (1991).

4. Vitousek, P. M. & Farrington, H. Biogeochemistry 37, 63–75 (1997).

5. Güsewell, S. N : P ratios in terrestrial plants: variation and functional significance. New Phytol. 164, 243–266 (2004).

6. Elser, J. J. et al. Global analysis of nitrogen and phosphorus limitation of primary producers in freshwater, marine and terrestrial ecosystems. Ecol. Lett. 10, 1135–1142 (2007).

7. Fisher, J. B., Badgley, G. & Blyth, E. Global nutrient limitation in terrestrial vegetation. Global Biochem. Cy. 26, GB3007 (2012).

8. Fay, P. A. et al. Grassland productivity limited by multiple nutrients. Nat Plants 1, 15080 (2015).

9. Chapin, F. S. I., Vitousek, P. M. & Van Cleve, K. The nature of nutrient limitation in plant communities. Am. Nat. 48–58 (1986).

10. von Liebig, J. Die organische Chemie in ihrer Anwendung auf Agricultur und Physiologie (F. Vieweg, 1841).

11. Van Sundert, K. et al. Towards comparable assessment of the soil nutrient status across scales-Review and development of nutrient metrics. Glob. Chang. Biol. 26, 392–409 (2020).

12. Koerselman, W. & Meuleman, A. F. M. The vegetation N:P ratio: A new tool to detect the nature of nutrient limitation. J. Appl. Ecol. 33, 1441 (1996).

13. Olde Venterink, H., Wassen, M. J., Verkroost, A. W. M. & De Ruiter, P. C. Species richness– productivity patterns differ between n-, p-, and k-limited wetlands. Ecology 84, 2191–2199 (2003).

14. Güsewell, S. & Koerselman, W. Variation in nitrogen and phosphorus concentrations of wetland plants. Perspect. Plant Ecol. Evol. Syst. 5, 37–61 (2002).

15. Tessier, J. T. & Raynal, D. J. Use of nitrogen to phosphorus ratios in plant tissue as an indicator of nutrient limitation and nitrogen saturation. J. Appl. Ecol. 40, 523–534 (2003).

16. Boeye, D., Verhagen, B., Van Haesebroeck, V. & Verheyen, R. F. Nutrient limitation in species-rich lowland fens. J. Veg. Sci. 8, 415–424 (1997).

17. Bedford, B. L., Walbridge, M. R. & Aldous, A. Patterns in nutrient availability and plant diversity of temperate north American wetlands. Ecology 80, 2151 (1999).

18. Wassen, M. J., Venterink, H. O., Lapshina, E. D. & Tanneberger, F. Endangered plants persist under phosphorus limitation. Nature 437, 547–550 (2005).

19. Ceulemans, T., Merckx, R., Hens, M. & Honnay, O. Plant species loss from European semi-natural grasslands following nutrient enrichment – is it nitrogen or is it phosphorus? Glob. Ecol. Biogeogr. 22, 73–82 (2013).

20. Palpurina, S. et al. The type of nutrient limitation affects the plant species richness–productivity relationship: Evidence from dry grasslands across Eurasia. J. Ecol. 107, 1038–1050 (2019).

21. Reich, P. B. & Oleksyn, J. Global patterns of plant leaf N and P in relation to temperature and latitude. Proc. Natl. Acad. Sci. U. S. A. 101, 11001–11006 (2004).

22. Houlton, B. Z., Wang, Y.-P., Vitousek, P. M. & Field, C. B. A unifying framework for dinitrogen fixation in the terrestrial biosphere. Nature 454, 327–330 (2008).

23. He, N. et al. Global patterns of nitrogen saturation in forests. Research Square (2023) doi:10.21203/rs.3.rs-3559857/v1.

24. Vitousek, P. M. & Reiners, W. A. Ecosystem succession and nutrient retention: A hypothesis. Bioscience 25, 376–381 (1975).

25. Batjes, N. H. Total carbon and nitrogen in the soils of the world. Eur. J. Soil Sci. 47, 151–163 (1996).

26. Van Duren, I. C. & Pegtel, D. M. Plant Soil 220, 35–47 (2000).

27. Rousk, J., Brookes, P. C. & Bååth, E. Contrasting soil pH effects on fungal and bacterial growth suggest functional redundancy in carbon mineralization. Appl. Environ. Microbiol. 75, 1589–1596 (2009).

28. Aerts, R., Wallen, B. & Malmer, N. Growth-limiting nutrients in sphagnum-dominated bogs subject to low and high atmospheric nitrogen supply. J. Ecol. 80, 131 (1992).

29. Aerts, R. & Chapin, F. S. C. The Mineral Nutrition of Wild Plants Revisited: A Re-Evaluation of Processes and Patterns. Adv. Ecol. Res. 30, 1–67 (2000).

30. Devau, N., Cadre, E. L., Hinsinger, P., Jaillard, B. & Gérard, F. Soil pH controls the environmental availability of phosphorus: Experimental and mechanistic modelling approaches. Appl. Geochem. 24, 2163–2174 (2009).

31. Chen, Y. et al. Multi-elemental stoichiometric ratios of atmospheric wet deposition in Chinese terrestrial ecosystems. Environ. Res. 245, 117987 (2024).

32. Walker, T. W. & Syers, J. K. The fate of phosphorus during pedogenesis. Geoderma 15, 1–19 (1976).

33. Giesler, R., Högberg, M., Högberg, P. Soil chemistry and plants in Fennoscandian boreal forest as exemplified by a local gradient. Ecology 79, 119–37 (1998).

34. Arai, Y. & Sparks, D. L. Phosphate reaction dynamics in soils and soil components: A multiscale approach. in Advances in Agronomy 135–179 (Elsevier, 2007).

35. Tunesi, S., Poggi, V. & Gessa, C. Nutr. Cycling Agroecosyst. 53, 219–227 (1999).

36. Devau, N., Hinsinger, P., Le Cadre, E., Colomb, B. & Gérard, F. Fertilization and pH effects on processes and mechanisms controlling dissolved inorganic phosphorus in soils. Geochim. Cosmochim. Acta 75, 2980–2996 (2011).

37. Weng, L., Van Riemsdijk, W. H. & Hiemstra, T. Factors controlling phosphate interaction with iron oxides. J. Environ. Qual. 41, 628–635 (2012).

38. Verhoeven, J. T., Koerselman, W. & Meuleman, A. F. Nitrogen- or phosphorus-limited growth in herbaceous, wet vegetation: relations with atmospheric inputs and management regimes. Trends Ecol. Evol. 11, 494–497 (1996).

39. Manu, R. et al. Response of tropical forest productivity to seasonal drought mediated by potassium and phosphorus availability. Nat. Geosci. 17, 524–531 (2024).

40. Manning, D. A. C. Mineral sources of potassium for plant nutrition. A review. Agron. Sustain. Dev. 30, 281–294 (2010).

41. Van Sundert, K. et al. Post-drought rewetting triggers substantial K release and shifts in leaf stoichiometry in managed and abandoned mountain grasslands. Plant Soil 448, 353–368 (2020).

42. Tinker, P. B. & Nye, P. H. Solute Movement in the Rhizosphere. (Oxford University Press, 2000).

43. Hoosbeek, M. R., Van Breemen, N., Vasander, H., Buttler, A. & Berendse, F. Potassium limits potential growth of bog vegetation under elevated atmospheric CO2 and N deposition. Glob. Chang. Biol. 8, 1130–1138 (2002).

44. Kotowski, W. & van Diggelen, R. Light as an environmental filter in fen vegetation. J. Veg. Sci. 15, 583–594 (2004).

45. LeBauer, D. S. & Treseder, K. K. Nitrogen limitation of net primary productivity in terrestrial ecosystems is globally distributed. Ecology 89, 371–379 (2008).

46. Skiba, U. et al. Biosphere–atmosphere exchange of reactive nitrogen and greenhouse gases at the NitroEurope core flux measurement sites: Measurement strategy and first data sets. Agric. Ecosyst. Environ. 133, 139–149 (2009).

47. Wardle, D. A., Walker, L. R. & Bardgett, R. D. Ecosystem properties and forest decline in contrasting long-term chronosequences. Science 305, 509–513 (2004).

48. Lammerts, E. J., Pegtel, D. M., Grootjans, A. P. & van der Veen, A. Nutrient limitation and vegetation changes in a coastal dune slack. J. Veg. Sci. 10, 111–122 (1999).

49. Berendse, F. Organic matter accumulation and nitrogen mineralization during secondary succession in heathland ecosystems. J. Ecol. 78, 197–215 (1990).

50. Roem, W. J. & Berendse, F. Soil acidity and nutrient supply ratio as possible factors determining changes in plant species diversity in grassland and heathland communities. Biol. Conserv. 92, 151–161 (2000).

51. Van Sundert, K., Horemans, J. A., Stendahl, J. & Vicca, S. The influence of soil properties and nutrients on conifer forest growth in Sweden, and the first steps in developing a nutrient availability metric. Biogeosciences 15, 3475–3496 (2018).

52. Chen, B. et al. A meta-analysis highlights globally widespread potassium limitation in terrestrial ecosystems. New Phytol. 241, 154–165 (2024).

53. de Mars, H., Wassen, M. J. & Peeters, W. H. M. The effect of drainage and management on peat chemistry and nutrient deficiency in the former Jegrznia-floodplain (NE-Poland). Vegetatio 126, 59–72 (1996).

54. Radujković, D. et al. Soil properties as key predictors of global grassland production: Have we overlooked micronutrients? Ecol. Lett. 24, 2713–2725 (2021).

55. Van Sundert, K. et al. When things get MESI: The Manipulation Experiments Synthesis Initiative-A coordinated effort to synthesize terrestrial global change experiments. Glob. Chang. Biol. 29, 1922–1938 (2023).

56. Gérard, F. Clay minerals, iron/aluminum oxides, and their contribution to phosphate sorption in soils — A myth revisited. Geoderma 262, 213–226 (2016).

57. Falk, K. et al. Molinia caerulea responses to N and P fertilisation in a dry heathland ecosystem (NW-Germany). Plant Ecol. 209, 47–56 (2010).

58. Roem, W. J., Klees, H. & Berendse, F. Effects of nutrient addition and acidification on plant species diversity and seed germination in heathland. J. Appl. Ecol. 39, 937–948 (2002).

59. Frosini, S., Lardicci, C. & Balestri, E. Global change and response of coastal dune plants to the combined effects of increased sand accretion (burial) and nutrient availability. PLoS One 7, e47561 (2012).

60. Van Sundert, K. et al. Fertilized graminoids intensify negative drought effects on grassland productivity. Glob. Chang. Biol. 27, 2441–2457 (2021).

61. Xu, H. et al. Impact of nitrogen addition on plant-soil-enzyme C–N–P stoichiometry and microbial nutrient limitation. Soil Biol. Biochem. 170, 108714 (2022).

62. Ellenberg, H. Indicator values of plants in Central Europe. (1992).

63. Tichý, L. et al. Ellenberg-type indicator values for European vascular plant species. J. Veg. Sci. 34 (2023).

64. Dengler, J. & Dembicz, I. Should we estimate plant cover in percent or on ordinal scales? Vegetation Classification and Survey 4, 131–138 (2023).

65. Walinga, I., van Vark, W., Houba, V. J. G., van der Leep, J. J. Plant analysis procedures. Soil and Plant Analysis (Agricultural University Wageningen, 1989).

66. Tichý, L. JUICE, software for vegetation classification. J. Veg. Sci. 13, 451–453 (2002).

67. Berg, C. Common file of EIVs for vascular plants, mosses and lichens (Karl-Franzens-Universität Graz, 2011).

68. Hennekens, S. M. & Schaminée, J. H. J. Turboveg, a comprehensive database management system for vegetation data. J. Veg. Sci. 12, 589–591 (2001).

69. Houba, V. J. G., van der Lee, J. J., Novazamsky, I., Walinga, I. Soil Analysis Procedures, Soil and plant analysis (Agricultural University Wageningen, 1989).

70. Runhaar, J., Jalink, M. H., Hunneman, H., Witte, J. P. M., Hennekens, S. M. Ecologische vereisten habitattypen (Watercycle Research Institute, 2009).

71. Directorate-General for Environment of the European Commission. Interpretation manual of European Union habitats (European Commission, 2013).

72. Schaminée, J., Stortelder, A. H. F. & Dijk, E. De Vegetatie van Nederland (Opulus, 1995).

73. Oksanen J. et al. vegan: Community Ecology Package. R package version 2.6–8 (2024).

74. Manders, A. M. M. et al. Curriculum vitae of the LOTOS–EUROS (v2.0) chemistry transport model. Geosci. Model Dev. 10, 4145–4173 (2017).

75. Kuenen, J., Dellaert, S., Visschedijk, A., Jalkanen, J.-P., Super, I. & Denier van der Gon, H. CAMS-REG-v4: a state-of-the-art high-resolution European emission inventory for air quality modelling. *Earth Syst*. Sci. Data 14, 491–515 (2022).

76. Schneider, C., Pelzer, M., Toenges-Schuller, N., Nacken, M. & Niederau, A. ArcGIS basierte Lösung zur detaillierten, deutschlandweiten Verteilung (Gridding) nationaler Emissionsjahreswerte auf Basis des Inventars zur Emissionsberichterstattung: Forschungskennzahl (Umweltbundesamt, 2016).

77. Van Zanten, M. C., Sauter, F. J., Wichink Kruit, R. J., Van Jaarsveld, J. A. & Van Pul, W. A. J. Description of the DEPAC module. Dry deposition modeling with DEPAC GCN2010 (Netherlands National Institute for Public Health and the Environment, 2010).

78. Wichink Kruit, R. J., Schaap, M., Sauter, F. J., van Zanten, M. C. & van Pul, W. A. J. Modeling the distribution of ammonia across Europe including bi-directional surface–atmosphere exchange. Biogeosciences 9, 5261–5277 (2012).

79. Hamilton, N. E. & Ferry, M. ggtern: ternary diagrams using ggplot2. J. Stat. Softw. 87, 1–17 (2018).

80. R Core Team. R: A Language and Environment for Statistical Computing (R Foundation for Statistical Computing, 2024).

81. Kuznetsova, A., Brockhoff, P. B., Christensen, R. H. B. lmerTest: Tests in Linear Mixed Effects Models. J. Stat. Softw. 82, 1–26 (2017).

82. Christensen, R. Ordinal-Regression Models for Ordinal Data. R package version 2023 (2023).

83. Bivand, R. R packages for analyzing spatial data: A comparative case study with areal data. Geogr. Anal. 54, 488–518 (2022).

84. Ripley, B. Tree: Classification and Regression Trees. R package version 1.0–43 (2023).

85. Hartig, F. DHARMa: Residual Diagnostics for Hierarchical (Multi-Level / Mixed) Regression. R package version 0.4.6 (2022).

86. Stoffel, M. A., Nakagawa, S. & Schielzeth, H. partR2: Partitioning R^2^ in generalized linear mixed models. bioRxiv (2020) doi:10.1101/2020.07.26.221168.

87. Lenth, R. V. Least-Squares Means: TheRPackagelsmeans. J. Stat. Softw. 69, (2016).

