## Supplementary text and figures for "Phosphorus availability as the primary determinant of nutrient limitation in temperate biodiverse herbaceous vegetation"

#### Text S1. Supplementary results.

**Characterization of habitat types.** Each of the 386 sampled communities was assigned a habitat type based on species' presence and ecology[72]. Six communities were categorized as calcareous springs, 99 as fens, 26 as bogs, 19 were wet dune slacks, 32 fen meadows, 18 *Nardus* grasslands, 51 wet heathlands, 14 dry calcareous grasslands, 11 dry dunes, and 11 dry heathlands. 99 plant communities were clearly in a degraded, species-poorer state or could not be assigned to one of these ten habitat types that fit in the European Natura 2000 classification[71], and were categorized more broadly as either wet eutrophic marshes ($n$ = 13), moist grasslands ($n$ = 65) or dry grasslands ($n$ = 21).

The ecological variables represented long gradients across the entire dataset. The Ellenberg indicator value (EIV) for soil moisture varied between 3.00 and 10.00. The habitat types differed in their position along this soil moisture axis ($F_{12,121}$ = 61.43, $P$ < 0.001; Extended Data Fig. 1a). Based on the vegetation present, calcareous springs, wet eutrophic marshes, fens and bogs were characterized as "wet" (Extended Data Table 1), showing little overlap with the next "moist" category encompassing wet dune slacks, fen meadows, *Nardus* grasslands, moist grasslands and wet heathlands. The "dry" habitats calcareous grasslands, dunes, dry grasslands and heathlands significantly differed in their moisture EIV from the other habitat types, except for some overlap between dry heathlands and the driest among *Nardus* grasslands, which varied substantially in their moisture EIV.

Soil pH also varied substantially with values measured *in situ* in the field or in extracts with water between 2.6 and 8.6, and pH measured in KCl solution ranging between 2.3 and 7.6. pH conditions thus varied from acidic ($pH_{water}$ < 4.5 or $pH_{KCl}$ < 3.5) to alkaline ($pH_{water}$ > 7.5 or $pH_{KCl}$ > 7.5), with many of the communities growing in intermediate conditions of moderate (4.5 < $pH_{water}$ < 5.5 or 3.5 < $pH_{KCl}$ < 4.8), weak (5.5 < $pH_{water}$ < 6.5 or 4.8 < $pH_{KCl}$ < 6.1), or circumneutral (6.5 < $pH_{water}$ < 7.5 or 6.1 < $pH_{KCl}$ < 7.5) acidity[70]. Following this classification system that integrates $pH_{water}$ and $pH_{KCl}$ into one single indicator, habitat types clearly differed in soil pH ($\chi^2_{12}$ = 176.46, $P$ < 0.001). Bogs, wet and dry heathlands could be categorized as clearly acidic habitat types, exhibiting significant overlap only with some *Nardus* grasslands. The latter fell into a broad category with generally weakly to moderately acidic soils, together with fens, fen meadows, moist grasslands, dry dunes and dry grasslands. Alkaline to circumneutral conditions occurred predominantly in calcareous springs, wet eutrophic marshes, wet dune slacks and dry calcareous grasslands in our dataset (Extended Data Fig. 1b; Extended Data Table 1).

**Nutrient stoichiometry among habitat types.** Elemental stoichiometries varied substantially. N:P ratios ranged between 3.3 and 47.2, N:K between 0.4 and 8.9, and K:P between 0.8 and 30.5. These ratios differed significantly amongst habitats (N:P: $F_{12,137}$ = 16.07, $P$ < 0.001; N:K: $F_{12,138}$ = 10.50, $P$ < 0.001; K:P: $F_{12,133}$ = 6.56, $P$ < 0.001), with bogs and wet heathlands showing an overall N:P ratio significantly greater than the critical value of 14.5[13], whereas dry dunes, wet eutrophic marshes, and moist and dry grasslands exhibited on average ratios significantly less than this critical value (Fig. 2a). N:K ratios of dry heathlands were on average significantly greater than the critical value of 2.1, and for wet heathlands, bogs and *Nardus* grasslands no significant deviation occurred from this critical ratio. All other habitat types had N:K ratios significantly lower than 2.1 (Extended Data Fig. 2a). Ratios of K to P were for all habitat types on average significantly or near significantly greater than the critical value of 3.4 (Extended Data Fig. 2b), with K:P of calcareous springs and dry calcareous grasslands particularly high (> 15). K:P of moist grasslands and dry heathlands was lowest (< 6, on average). However, in individual plant communities, K:P ratios below 3.4 did occur occasionally in some dry heathlands ($n$ = 1), fen meadows ($n$ = 1) and *Nardus* grasslands ($n$ = 2), somewhat more in dry grasslands ($n$ = 5) and fens ($n$ = 6), and several times in moist grasslands ($n$ = 24).

### Supplementary figures

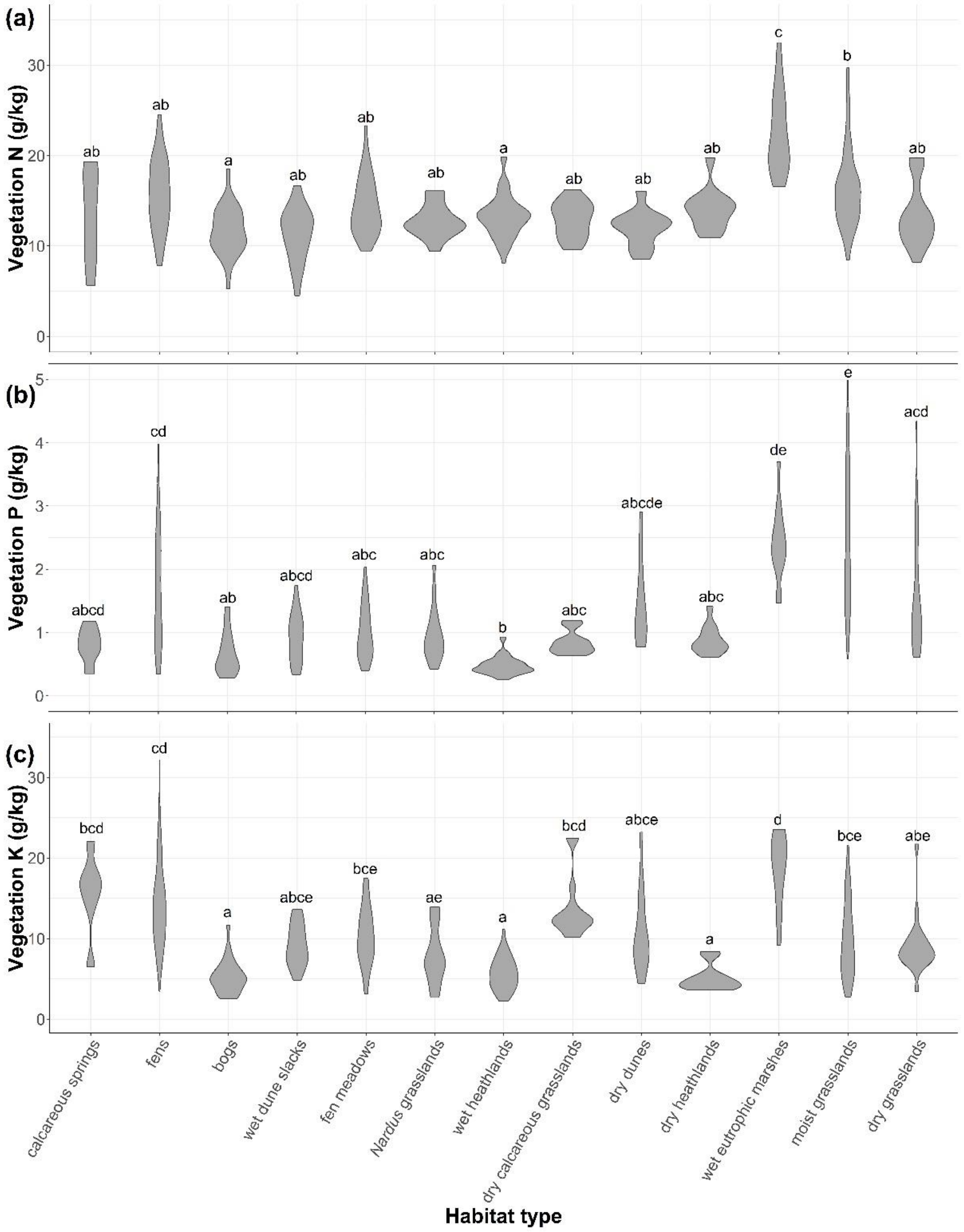


65 **Fig. S1 | Vegetation N, P and K concentrations across habitat types.** Values significantly different among habitat types were indicated trough different letters.

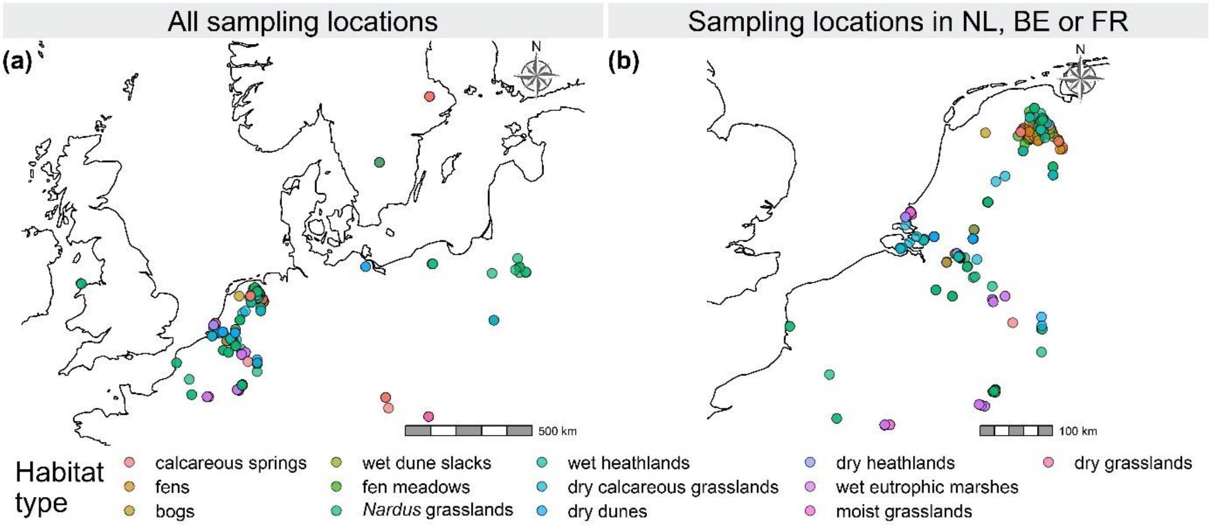


**Fig. S2 | Overview of sampling locations in temperate northwest and central Europe.**

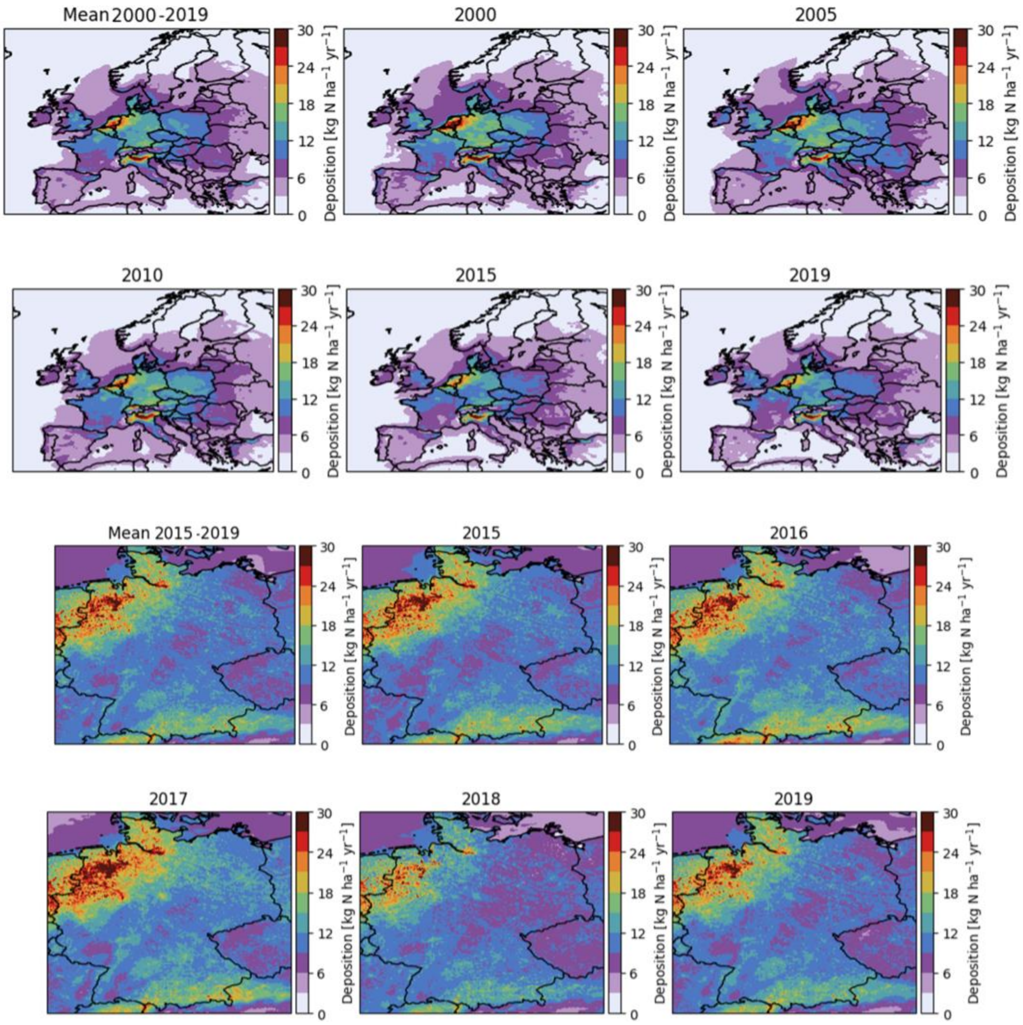


**Fig. S3 | Spatiotemporal variation in modeled N deposition.** Model output at 25 x 25 km² in Europe (2000-2019 mean & individual years 2000, 2005, 2010, 2015, 2019, top six panels) and 2 x 2 km² resolution in northwest Europe (2015-2019 mean & individual years, bottom six panels).
